# Structural conversion of α-synuclein at the mitochondria induces neuronal toxicity

**DOI:** 10.1101/2022.06.07.494932

**Authors:** Minee L. Choi, Alexandre Chappard, Bhanu P. Singh, Catherine Maclachlan, Margarida Rodrigues, Evgenia Fedotova, Alexey V. Berezhnov, Suman De, Chris Peddie, Dilan Athauda, Gurvir S. Virdi, Weijia Zhang, James R. Evans, Anna Wernick, Zeinab Shadman Zanjani, Plamena R. Angelova, Noemi Esteras, Andrey Vinikurov, Katie Morris, Kiani Jeacock, Laura Tosatto, Daniel Little, Paul Gissen, David J. Clarke, Tilo Kunath, Lucy Collinson, David Klenerman, Andrey Y. Abramov, Mathew H. Horrocks, Sonia Gandhi

## Abstract

Aggregation of α-Synuclein (α-Syn) drives Parkinson’s disease, although the initial stages of self-assembly and structural conversion have not been captured inside neurons. We track the intracellular conformational states of α-Syn utilizing a single-molecule FRET biosensor, and show that α-Syn converts from its monomeric state to form two distinct oligomeric states in neurons in a concentration dependent, and sequence specific manner. 3D FRET-CLEM reveals the structural organization, and location of aggregation hotspots inside the cell. Notably multiple intracellular seeding events occur preferentially on membrane surfaces, especially mitochondrial membranes. The mitochondrial lipid, cardiolipin triggers rapid oligomerization of A53T α-Syn, and cardiolipin is sequestered within aggregating lipid-protein complexes. Mitochondrial aggregates impair complex I activity and increase mitochondrial ROS generation, which accelerates the oligomerization of A53T α-Syn, and ultimately causes permeabilization of mitochondrial membranes, and cell death. Patient iPSC derived neurons harboring A53T mutations exhibit accelerated oligomerization that is dependent on mitochondrial ROS, early mitochondrial permeabilization and neuronal death. Our study highlights a mechanism of de novo oligomerization at the mitochondria and its induction of neuronal toxicity.

## Introduction

The presence of aggregates rich in alpha-synuclein (α-Syn) is a fundamental hallmark of synucleinopathies, a subset of neurodegenerative diseases that encompass Parkinson’s disease (PD), Dementia with Lewy bodies (DLB) and Multiple system atrophy (MSA) [1–4]. The α-Syn protein plays a central role in the pathogenesis of these diseases. Supportive of this are genetic studies confirming that mutations and multiplications in the SNCA gene on chromosome 4q21-23, which encodes for α-Syn, cause early-onset familial PD [4–7], with widespread deposition of α-Syn aggregates in the brain. α-Syn is an intrinsically disordered protein, and during pathology, undergoes a conformational change from its native state to β-sheet-rich structures including oligomers, protofibrils, and insoluble fibrils that finally accumulate in Lewy bodies (reviewed here [8]). Although it is the insoluble end-stage species of protein aggregation that has traditionally defined disease, it is the soluble, intermediate species of oligomers formed throughout aggregation which are markedly toxic, inducing aberrant calcium signaling, generating reactive oxygen species, mitochondrial dysfunction and neuronal cell death compared to other forms [9–13]. The early dynamic processes involved in the formation and toxicity of α-Syn oligomers are difficult to investigate using traditional methods due to the intrinsically transient, heterogeneous and low abundant nature of oligomers (reviewed here [14]). To overcome this, an array of single-molecule fluorescence methods has been developed to observe protein interactions and conformation, which broadly rely on the detection of transfer of excitation energy between two fluorophores if sufficiently close (reviewed here [15]). By employing single-molecule Förster resonance energy transfer (smFRET), we were previously able to track α-Syn aggregation *in vitro*, distinguish different structural groups of oligomers formed during aggregation, and compare the kinetics of oligomer formation from patients with familial PD mutations, demonstrating A53T α-Syn has a higher propensity to form cytotoxic oligomers compared to other mutations [16, 17]. However it has not yet been possible to capture and characterize the early dynamic process of aggregation inside the native human environment, and determine its effects on cellular homeostasis.

In this study, we integrated sensitive biophysical approaches including a FRET biosensor, single-molecule FRET analyses, and FRET-Correlative light and electron microscopy (CLEM) imaging, to precisely track live the kinetics, triggers, and location of α-Syn aggregation inside rodent and human neurons. This allowed us to visualize the maturing aggregate in its native location, and led us to investigate how protein-lipid interactions may trigger aggregation and alter the kinetics of aggregation. Applying high resolution methods to resolve this key process revealed a mechanism by which mutant α-Syn aggregates in mitochondria, and alters mitochondrial bioenergetics, and ultimately leads to toxicity.

## Results

### FRET biosensor can detect intracellular oligomerization and characterize the kinetics

Fluorescently labeled α-Syn was used to measure uptake of α-Syn by neurons, and its subsequent aggregation state by intracellular FRET. Two populations of α-Syn, one labelled with Alexa Fluor 488 (AF488) and the second with Alexa Fluor 594 (AF594) (Figure 1a) were mixed and added to neurons, and the intracellular accumulation of α-Syn was visualized by measuring the intensity of AF594 within the cells using direct excitation with 594 nm irradiation (Figure 1bi, ci & S1a). This direct excitation signal gives a measure of the total α-Syn added to the cells, regardless of its aggregation state. The formation of oligomers was visualized via the presence of signal from the acceptor fluorophore (AF549) after excitation of the donor fluorophore with 488 nm irradiation, which occurs as energy is non-radiatively transferred to from AF488 to AF594. We refer to this as the FRET signal, and it can only occur when the fluorophores are in close proximity (< 10 nm), as is the case within aggregates. We corrected for fluorescence cross-talk between the channels, direct excitation of the acceptor fluorophore and autofluorescence, and confirmed that cells treated with either AF488 or AF594 alone did not induce a FRET signal (Figure S1bi).

**Figure 1.**
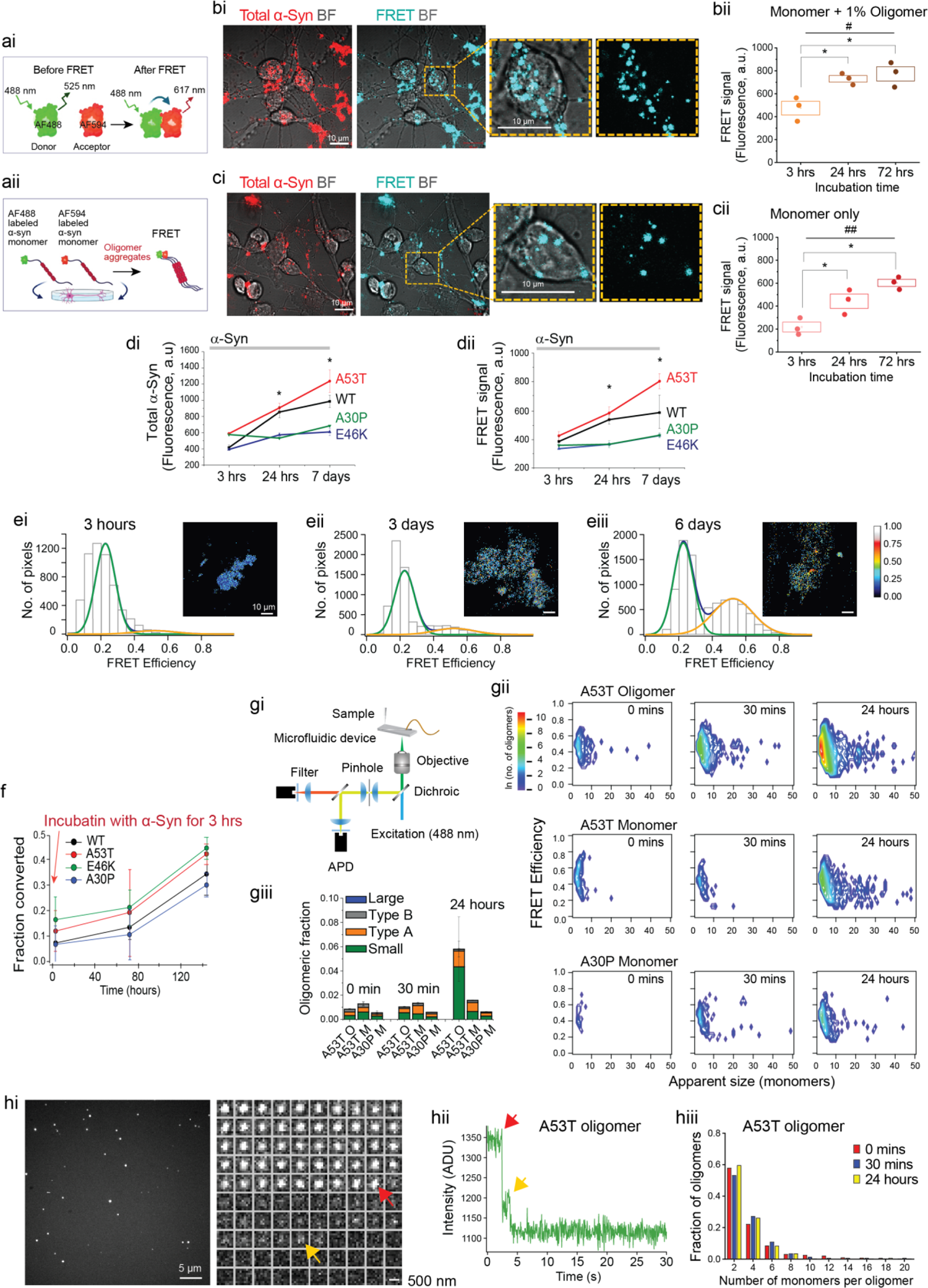
FRET sensor detects rapid intracellular oligomerization of A53T α-Syn. **(ai)** Schematic illustration demonstrating how the FRET sensor detects aggregation. Upon forming oligomers by binding two monomers within 10 nm, energy is transferred from the donor fluorophore (AF488) to the acceptor fluorophore (AF594). **(aii)** α-Syn-AF488 and α-Syn-AF594 monomers are applied to cells and the FRET signal is detected. **(bi)** Representative FRET images after 72 hours incubation with oligomers. **(bii)** Application of 500 nM WT oligomeric α-syn has detectable FRET which increases over time (3 hrs vs 24 hrs: *p* = 0.032, 3 hrs vs 72 hrs: *p* = 0.003). **(ci)** Representative FRET images after 72 hours incubation with monomers. **(cii)** Application 500 nM monomeric α-Syn exhibits low FRET signal initially, followed by an increase in FRET over time (3 hrs vs 24 hrs: *p* = 0.031, 3 hrs vs 72 hrs: *p* = 0.016). **(di)** A53T monomer exhibits highest intracellular accumulation. **(dii)** A53T monomer exhibits highest intracellular FRET intensity over. **(ei – iii)** Structural conversion from low FRET efficient Type-A oligomers to high FRET efficient Type-B oligomers over 6 days. **(f)** Short incubation with WT and mutant monomers results in increase in FRET efficiency over time. **(gi)** Single-molecule confocal microscopy under conditions of fast-flow used to analyze cell lysates. **(gii)** Two-dimensional contour plots of approximate oligomer size and FRET efficiency after application of A53T oligomer, monomer, and A30P monomer. Both the number of events and the size of the oligomers increase over time in all cases. **(giii)** Number and type of oligomeric events in cell lysates from A53T oligomer, A53T monomer, and A30P monomer treated cells. **(hi)** Photobleaching step analysis for A53T oligomer treated cells. **(hii)** Step-fit example of a single A53T oligomer (24 hours) trace. **(hiii)** Photobleaching trace data was filtered and then fitted to show indicative photobleaching steps. Each step indicates photobleaching of a single fluorophore, from which oligomer size can be estimated. *Note*. Data are represented as mean ± SEM; n = 2 – 7 number of wells. Total number of cells = 46 – 214. All experiments were independently repeated 2 – 3 times. *p < 0.05, **p < 0.001, ***p < 0.0001. See also Figure S1 & 2.

We applied 500 nM oligomer sample (containing ∼1 % oligomer, ∼99 % monomer) formed from the AF488-α-Syn and AF594-α-Syn, to primary neurons and measured intracellular FRET using live-cell imaging at different time points. Uptake of α-Syn resulted in both a direct excitation signal (total α-Syn) and FRET signal (oligomer formation), which both increased in intensity over time (474.3 ± 60.1, 730.3 ± 28.7 and 774.0 ± 62.1 a.u. at 3 hours, 24 hours and 72 hours incubation respectively), reflecting the continual uptake of total α-Syn, and for oligomers to seed aggregation inside cells (Figure 1bi & ii). Aggregation of α-Syn is a concentration dependent process. Application of an initial concentration of 5 – 50 nM α-Syn (containing 0.05 – 0.5 nM oligomer) induced intracellular oligomerization after 24 hours incubation (Figure S1ci), whilst 100 – 500 nM α-Syn (containing 1 - 5 nM oligomer) induces oligomerization within 3 hours (Figure S1cii). In order to verify that the intracellular FRET signal is a biosensor of oligomer formation, we confirmed FRET co-localization with an ATTO425 labelled aptamer specific to oligomeric aggregates (0.67 ± 0.07 ratio, Figure S1E) [18, 19], and areas of co-localization between a conformation specific antibody and the FRET signal (0.28 ± 0.05 ratio, Figure S1di & ii) [12, 19].

Next we tested whether application of α-Syn monomers alone (in the absence of any seeds) can oligomerize by applying 500 nM AF488-α-Syn and AF594-α-Syn monomers to cells. Interestingly, following an initial lag phase, the FRET intensity increased over time (217.6 ± 42.3, 442.6 ± 62.3 and 602.6 ± 31.9 a.u. at 3 hours, 24 hours and 72 hours incubation respectively. Figure 1ci & ii). Next, we treated cells with 500 nM AF488-α-Syn and AF594-α-Syn monomers of A53T, A30P, E46K or wild-type (WT), and measured both the direct excitation signal (total α-Syn) and intracellular FRET (oligomer formation) at different time-points. The uptake increased for the WT and all mutants over time (Figure 1di). The FRET signal also increased for WT and all mutants over time, showing a time-dependent increase in aggregation (Figure 1dii). Of note, A53T showed the greatest increase in both direct excitation signal and intracellular FRET signal over time indicating increased uptake, and oligomerization compared to the other mutations (A53T: 1236 ± 140, WT: 987 ± 75, A30P: 609 ± 43, E46K: 684 ± 4 a.u. at 72 hours).

This experimental paradigm can detect different conformational states (monomer vs oligomer) inside cells, and demonstrates that intracellular monomer begins to self-assemble to oligomeric states in a time dependent, concentration dependent, and sequence specific manner.

### Structural conversion measured by FRET efficiency

Single-molecule FRET measurements can identify the *in vitro* structural conversion from less toxic, loosely associated “Type-A” oligomers into toxic, proteinase-K resistant, beta-sheet rich “Type-B” oligomers [20]. Taking advantage of the intracellular FRET signal detected here, we determined whether such a conversion could occur within neurons. To remove the effect of continued uptake of α-Syn, we replaced the media containing AF488 and AF594 labelled α-Syn with fresh media after three hours of incubation. We then measured the FRET efficiency (E) of the aggregates within the cells according to Equation 1.

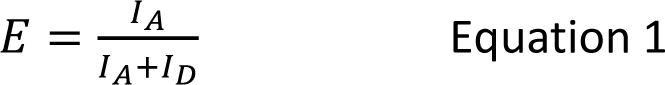

*W*here *I_A_* and *I_D_* are the intensities of the acceptor and donor fluorophores, respectively, with only 488 nm excitation. In this case *I_A_* is detectable when FRET occurs.

FRET efficiency histograms were generated from the intracellular aggregates following incubation with WT 500 nM α-Syn monomer and the histograms were fitted to two Gaussian distributions and integrated to obtain the fraction of converted oligomers (Figure 1e). At early time-points (3 hours and 3 days) only low FRET efficiency Type-A oligomers are present. After 6 days, however, a second high FRET efficiency peak appears, which corresponds to the Type-B oligomers previously identified *in vitro*. Assembly from monomer to Type-A oligomer occurs rapidly with a short lag phase < 3 hours, and conversion from Type-A to Type-B oligomers occurs over several days.

To determine the effect of the mutants to structurally convert, we removed the monomer from the media after 3 hours of incubation and measured FRET efficiency at a series of time-points. FRET efficiency, reflecting structural conversion, increased over the time for WT and mutants, and in the absence of continued uptake from the extracellular media, there was no difference in the kinetics of structural conversion between WT and mutations. (Figure 1f).

### A53T exhibits accelerated intracellular oligomerization with reduction of lag phase

To characterize the effect of the A53T mutation on aggregation (when continuously present), we applied AF488-α-Syn and AF594-α-Syn A53T or A30P, in monomeric or pre formed oligomeric state to cells, collected cell media and lysates at different time points, and used single-molecule confocal microscopy to measure the FRET efficiencies of individual oligomers, described in Figure 1gi [20]. Similar to the aggregation process *in vitro*, we were able to detect both low- and high-FRET oligomers with a range of different sizes as shown in the two-dimensional contour plots (Figure 1gii). Fitting of the resultant FRET efficiency histograms (Figure S2b), allowed the small, Type-A, Type-B and large oligomers to be quantified (Figure 1giii). As a positive control, A53T oligomer treated cells (1 % oligomer, 99 % monomer) exhibited rapid assembly to form oligomeric species of increasing size over 24 hours. A53T monomer treated cells exhibited rapid assembly into small oligomeric species at 0 – 30 min, which increased in size over 24 hours. Self-assembly and the early steps of aggregation were also observed in A30P treated cells, although less than A53T treated cells.

Single-molecule confocal microscopy only allows the approximate size distribution of the oligomers to be measured, as the total fluorescence intensity is dependent on a number of other factors in addition to the total number of fluorophores present. We therefore used total internal reflection fluorescence microscopy (TIRFM) to image the oligomers (Figure 1hi) [21–23]. As each α-Syn monomer carries a single dye molecule, and upon excitation, each label within an immobilized oligomer photobleaches, there is a step-wised decrease in intensity. By counting the number of these photobleaching steps, the number of monomers per oligomer can be determined (Figure 1hi & ii). These analyses were performed on lysates after the addition of A53T or A30P monomer and oligomer, and histograms showing the number of monomers-per-oligomer were generated (Figure 1hiii and S2c). These size distributions correlated well with those measured using single-molecule confocal microscopy (Figure 1hiii); approximately half of the population of oligomers contained two monomers, whilst the rest contained 4 – 10 monomers. However, it was possible to detect oligomers with up to 20 monomer units. Due to simultaneous photobleaching of multiple fluorophores, larger oligomers could not be distinguished using this method.

In agreement, an α-Syn oligomer specific ELISA demonstrated higher intracellular α-Syn oligomeric concentration (14.0 ± 1.8 nM after 24 hours and 16.3 ± 0.7 nM after 3-day treatment) in A53T monomer treated cells compared to WT monomer treated cells (12.3 ± 0.6 nM after 1-day and 13.1 ± 1.3 nM after 3-day treatment) (Figure S2d).

Taken together, the intracellular FRET, smFRET and TIRF microscopy can detect the initial stages of self-assembly, oligomer formation, and structural conversion inside cells. These methods track only the fluorescent labelled recombinant α-Syn, rather than the aggregation state of the endogenous (rodent) α-syn. Aggregation inside cells is concentration and time dependent, in agreement with aggregation experiments performed in buffer. However, in contrast to the kinetics of the aggregation in buffer in test tubes, there is a markedly reduced lag phase inside cells. Finally, there is an increased rate of oligomerization for A53T compared to WT and other mutants, and that this is in part due to enhanced uptake, and therefore increased intracellular concentration of A53T.

### Oligomer formation occurs in multiple cell ‘hotspots’ at heterogeneous locations

To study the ultrastructural location of oligomer formation in cells, we combined FRET imaging of labelled α-Syn with serial section Electron microscopy (EM), and with Focussed Ion Beam milling combined with Scanning Electron Microscopy (FIB-SEM) at 5 nm voxel resolution, to perform 3D FRET-CLEM in hiPSC derived neurons. We applied AF488-α-Syn and AF594-α-Syn A53T monomer and oligomer to neurons, generated FRET intensity heatmaps of aggregate formation, and tracked the same cells through sample preparation for serial section EM, highlighting multiple areas within the cell corresponding to different FRET intensities. We then performed a time course of AF488-α-Syn and AF594-α-Syn A53T monomer in cells, measuring intracellular FRET at different time-points (3 hours, 24 hours and 7 days) (Figure 2ai). The FRET intensity heatmaps reveal that aggregates form when monomers come into close proximity at areas of high concentration (within 3 hours). By 24 hours, the aggregates mature, showing a high FRET intensity in the center surrounded by a rim of lower FRET (indicating fewer, and loosely packed aggregates). Finally, after 7 days, these structures mature further into large clusters of aggregates (Figure 2aii) with central core of high FRET and rim of lower FRET signal.

Integrating FRET heatmaps with the EM images, and aligning for nuclear marker only, allowed some integrated spatial information to be obtained. At 3 hours, and 24 hours, we demonstrate that FRET, combined with FIB-SEM, shows that early oligomer formation occurs in a range of localizations to regions of the cell that encompass nucleus, mitochondria, and cytosol (3 hours and 24 hours, Figure 2b & c). After incubation with A53T for 7 days, the FRET signal, combined with serial section EM, showed that the maturing aggregates (7 days, Figure 2d) expand to occupy regions of the cell that encompass several different organelles, as well as the cytoplasm of the cell body in between organelles.

**Figure 2.**
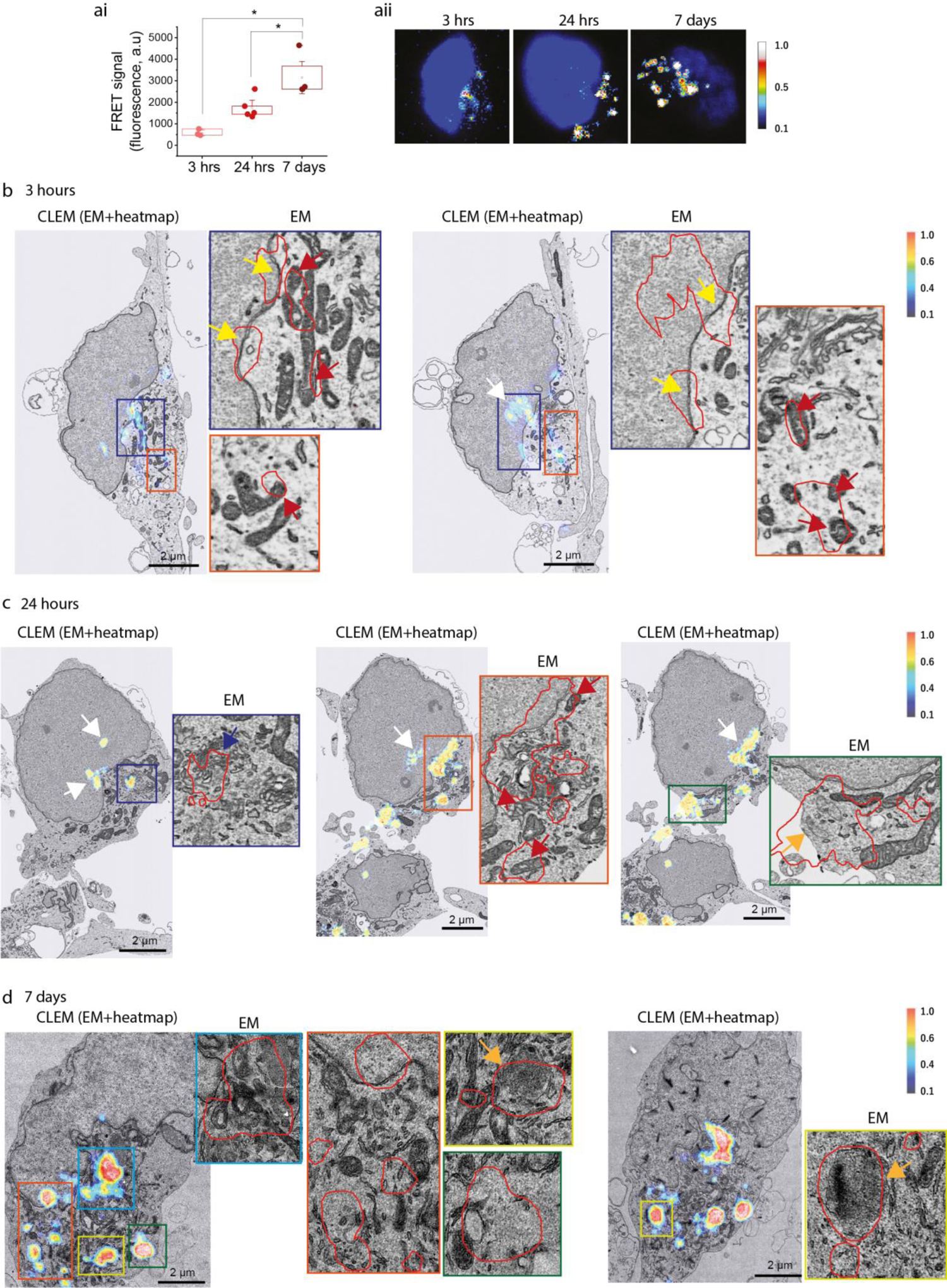
Oligomer formation occurs in multiple cell ‘hotspots’ at heterogeneous locations. **(ai)** Intracellular FRET intensity increases after application of 500 nM A53T monomer (3 hrs vs 7 days: *p* = 0.002, 24 hrs vs 7 days: *p* = 0.031) **(aii)** FRET intensity heat maps showing aggregate formation with high FRET signal intensity at the core, and low FRET intensity in the surrounding rim. **(b – d)** Application of equimolar concentration of A53T α-Syn-AF488 and A53T α-Syn-AF594 (total 500 nM), and images are obtained at three different time points, 3 hours, 24 hours and 7 days. Each panel is composed of a confocal image of CLEM (EM + FRET heat map) and zoom of EM alone. Colored arrows indicate aggregates at mitochondria (red), nucleus (white), membrane (yellow), Golgi apparatus (blue) and vesicles (orange). *Note*. Data are represented as mean ± SEM; n = 1 – 3 number of dishes. Total number of cells = 8 – 24. All experiments were independently repeated 2 – 3 times. *p < 0.05, **p < 0.001, ***p < 0.0001. See also Figure S3.

Refinements to the newly developed FRET-CLEM workflow would be required to understand the localization of oligomers at higher resolution. However, once an aggregation hotspot has formed, in a cellular space crowded with organelles, it will then mature in a stereotypical manner into an aggregate that contains highly ordered aggregates at the centre, and loosely ordered protein in the rim. The distribution of the early aggregation of A53T is clearly heterogenous, and there are multiple locations within the cell that act as a ‘seed’ for aggregation. This spatial information highlights how multiple seeding events in human cells may contribute to the reduced lag phase of aggregation that was observed using FRET.

### Cardiolipin triggers, and accelerates, the aggregation of A53T

Amongst the locations observed in the FRET-CLEM data were areas of the cell enriched in mitochondria and mitochondrial membranes. We then investigated whether the aggregation of A53T is promoted within mitochondria. Therefore, we performed an *in vitro* study of the kinetics of A53T and WT α-syn aggregation in the presence of cardiolipin. We used circular dichroism spectroscopy to demonstrate that α-syn binds cardiolipin, confirming that it adopts an alpha-helical conformation in the presence of cardiolipin liposomes (Figure S4a). Next we applied 10 μM α-syn to different lipids and found that phosphatidylserine-lipid (PS-lipid) vesicles promoted aggregation to fibrils, while phosphatidylcholine (PC-lipid) vesicle did not promote the aggregation of α-syn (Figure S4b). Addition of cardiolipin also triggered the aggregation of WT α-syn to form fibrils. Notably, application of 10 μM A53T monomer resulted in rapid A53T aggregation, with an absence of the lag phase in the presence of cardiolipin (Figure 3ai & ii). Conversely a long lag phase was observed with significant variation in the lag time for WT α-Syn (Figure 3aiii & iv). This implies that A53T monomers have higher propensity to aggregate in the presence of cardiolipin lipid vesicles than WT α-Syn. We then studied the morphology of the fibrils formed in the presence of cardiolipin by transmission electron microscopy (TEM). We found the morphological features of fibrils produced in the presence of cardiolipin are significantly different to those formed in the absence of cardiolipin, showing helical periodicity along the length of fibrils, and a greater number of protofilaments (Figure 3bi & ii). Thus, cardiolipin lipid vesicles may promote the lateral association of α-Syn protofilaments, which is consistent with the hierarchical self-assembly model of amyloid fibrils [24]. We next performed an aggregation assay with A53T α-Syn in the presence of 2 % biotinylated cardiolipin and subsequently imaged the lipid using Alexa Fluor 647 labelled streptavidin, and the fibrils using SAVE imaging on a TIRF microscope. This revealed that amyloid fibrils of A53T α-Syn are strongly co-localized with cardiolipin (84 ± 15 % coincident in comparison to the control (0.6 ± 1.2 %) without biotinylated lipid (Figure 3c). Interestingly, a previous study of A53T with phospholipid bilayers shows extraction and clustering of lipid molecule on the growing amyloid fibrils [25]. Our TIRFM data shows that the interaction of A53T α-Syn and lipid vesicles not only results in extraction of the lipid molecule cardiolipin, but furthermore the incorporation of cardiolipin within the amyloid fibrils. Our work suggests that cardiolipin rapidly triggers the aggregation of A53T, and is then incorporated into aggregating states of α-Syn which may further accelerate the aggregation process.

**Figure 3.**
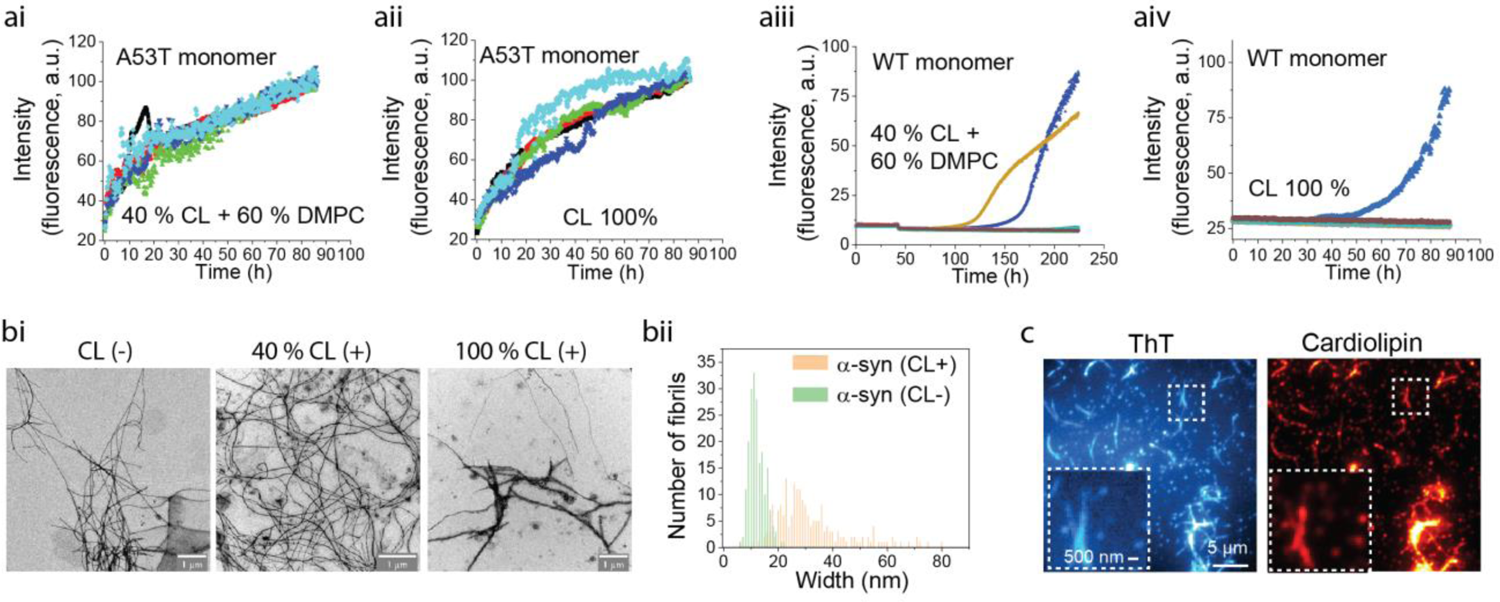
Cardiolipin triggers, and accelerates, the aggregation of A53T. **(ai – iv)** 50 µM A53T monomer led to a dramatically fast increase in ThT fluorescence in the presence of 40 % or 100 % cardiolipin as compared to WT monomer. **(bi)** TEM images show that in the presence of cardiolipin, fibrils of α-Syn have different morphology. A total 200 fibrils were analyzed for each group by Image J software [26]. **(bii)** Quantitative histogram of fibril width shows large distribution of width in the presence of cardiolipin, which is expected for a hierarchical self-assembly model of amyloid formation. **(c)** TIRFM analysis shows co-localization of cardiolipin and α-Syn fibrils (ThT positive). 0.5 µM α -Syn amyloid fibrils and in the presence of 4 µM cardiolipin vesicles (containing 2 mol % of biotinylated lipid) in the presence of 1 nM streptavidin-AF647 and 5 µµM ThT. Images were recorded for 50 frames from the red channel (AF647 emission) with 641 nm illumination, followed by green channel (ThT emission) with 405 nm illumination. *Note*. Data are represented as mean ± SEM; n = 3 independent experiments. See also Figure S4.

### A53T impairs mitochondrial bioenergetics and induces mitochondrial dysfunction

Next, we investigated the functional consequence of A53T α-Syn on mitochondrial function when applied exogenously to primary rodent neurons. A53T and WT α-Syn were applied to cells in their monomeric state, and single cell live confocal imaging was used to assess mitochondrial physiology. We measured the autofluorescence of reduced NADH which is a marker for cellular redox state, and mitochondrial electron transport chain function (reviewed here [27]). Application of 500 nM A53T induced an increase in NADH fluorescence, suggesting an impairment of Complex I function and accumulation of NADH similar to the effect of oligomeric α-Syn (Figure 4ai) [28]. Application of the same concentration of A53T monomer induced 21 % mitochondrial depolarization (Figure 4bi), measured using Rhodamine 123 fluorescence. NADH fluorescence could be partially restored by preincubation of cells with substrates for Complex I (Pyruvate, 5 mM) or Complex II (membrane permeable analogue of Succinate-di methyl succinate (DMsuccinate, 5 mM) reflecting that Complex I function, although impaired, could be rescued functionally (Figure 4aii), and that improvements in the respiratory chain function also drive restoration in the mitochondrial membrane potential (Δψ_m_), measured by Rhodamine123 (Figure 4bii). We also examined Δψ_m_ using the indicator tetramethylrhodamine methyl ester (TMRM). A53T reduced TMRM signal by 25 % after 30 min, whilst the TMRM signal was unchanged in WT treated cells (Figure 4c). We further investigated the cause of decreased Δψ_m_ by testing its sensitivity to Complex I and V inhibition. In healthy mitochondria, Δψ_m_ is maintained by Complex I – IV via electron transport chain function. WT treated cells maintain their Δψ_m_ predominantly through the action of complex I dependent respiration (exhibiting a 67 % reduction in Δψ_m_ following complex I inhibition). However A53T treated cells exhibit 37 % reduction of Δψ_m_ following complex V inhibition with oligomycin. (Figure 4di - iii). Therefore in cells exposed to A53T mutant, the Δψ_m_ cannot be maintained sufficiently through respiration, and must utilize Complex V (in reverse mode as an ATPase) to maintain it.

**Figure 4.**
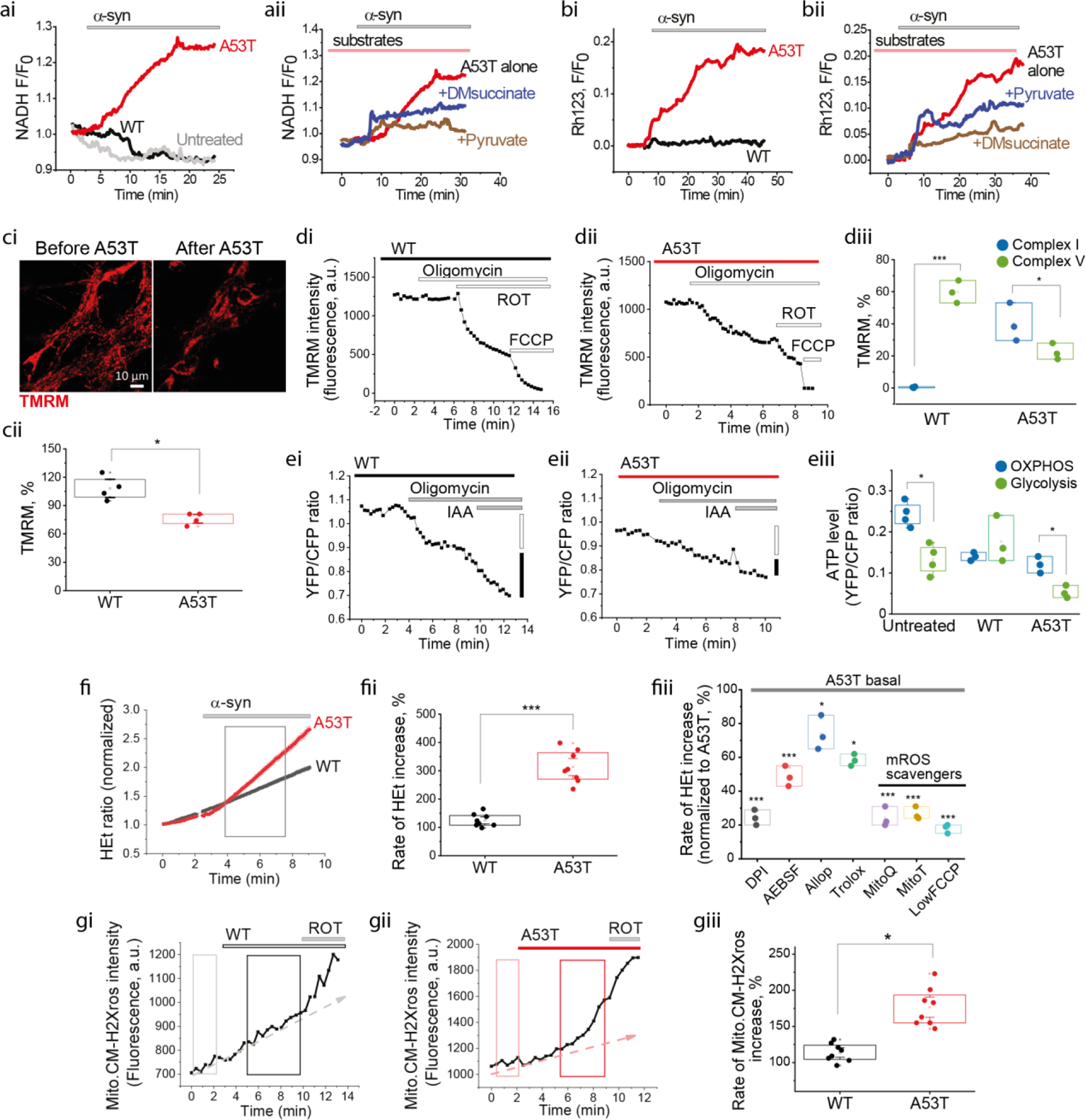
A53T impairs mitochondrial bioenergetics and induces mitochondrial dysfunction. **(ai)** Increase in NADH autofluorescence after application of 500 nM A53T α-syn (normalized to 1). **(aii)** The increase in NADH is prevented by a pre-incubation with Pyruvate and Succinate. **(bi)** A53T monomer depolarizes Δψ_m_ as measured by an increase in Rhodamine 123 fluorescence. **(bii)** The decreased Δψ_m_ is also reversed by pre-application of Pyruvate and Succinate. **(ci & ii)** Images demonstrating reduction in Δψ_m_ after 30 min incubation with A53T compared to WT (*p* = 0.021) and the quantitative histogram. **(di – iii)** Response of Δψ_m_ to Complex V inhibitor (Oligomycin: 2.4 μg/ml), Complex I inhibitor (Rotenone: ROT 5 μM), and mitochondrial uncoupler (FCCP: 1 μM). The basal fluorescence intensity was re-set at 1500 – 2500 a.u. **(ei – iii)** A53T reduces the total ATP production measured by FRET-ATP sensor, compared to WT treated (*p* = 0.03) or untreated cells (*p* = 0.026). **(fi & ii)** Superoxide is increased after application of A53T (*p* < 0.0001) but not WT. (fiii) Inhibition of A53T induced ROS by different inhibitors (0.5 µM DPI, 20 µM AEBSF, 20 µM Allop (Allopurinol), 100 µM Trolox, 0.1 µ MmitoQ, 0.1 µM MitoT (Mito-TEMPO) and 0.3 µM FCCP). **(gi – iii)** mROS production is increased by A53T (*p* = 0.012)*. Note*. 500 nM α-Syn monomer was applied for each experiment unless otherwise mentioned. Data are represented as mean ± SEM; n = 4 – 6 number of wells. Total number of cells = 215 – 481. All experiments were independently repeated 2 – 3 times. \**p* < 0.05, \*\**p* < 0.001, \*\*\**p* < 0.0001. See also Figure S5.

**Figure 5.**
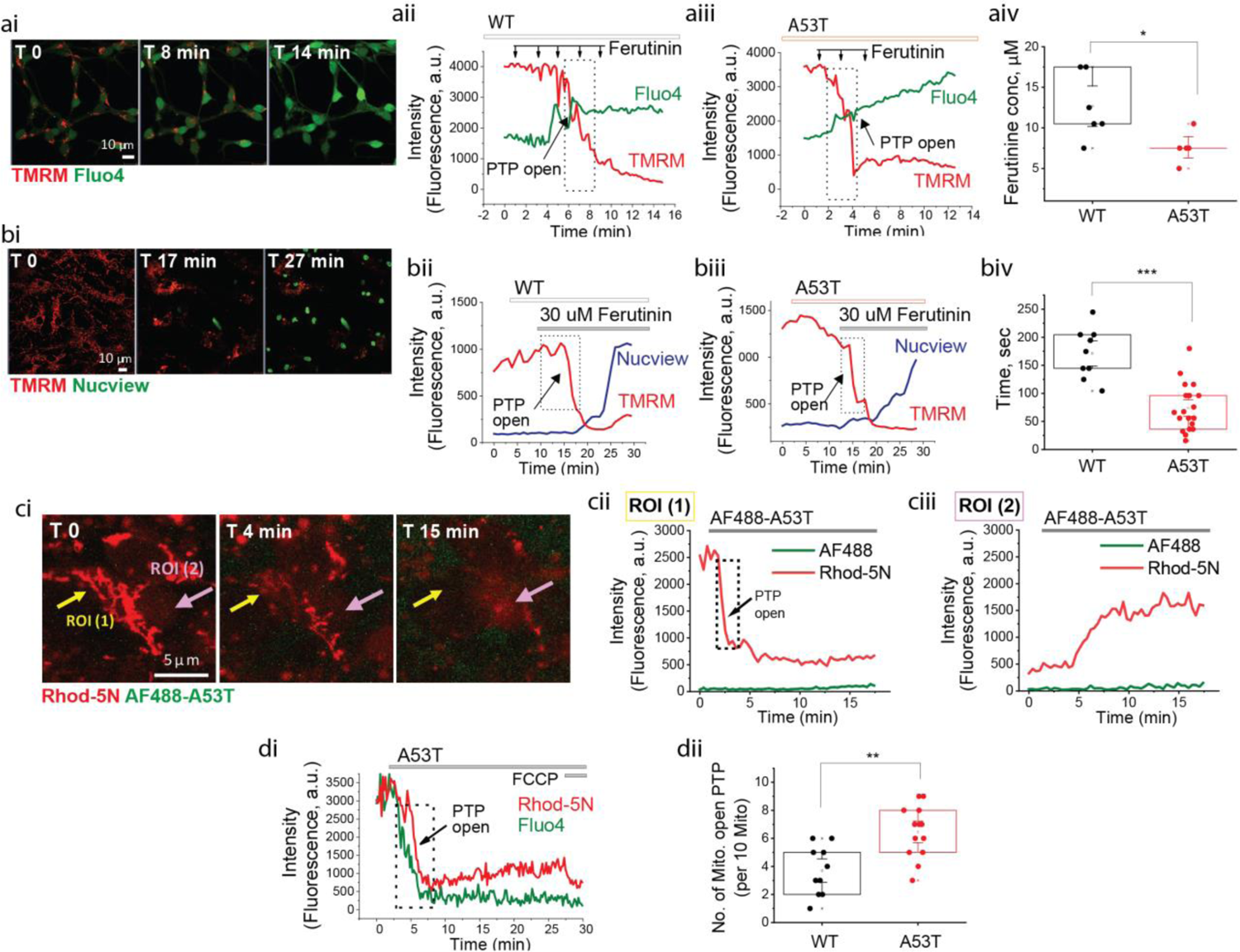
A53T induces mPTP opening. **(ai)** Representative time course images showing Δψ_m_ reduction (TMRM) is followed by an increase of cytoplasmic calcium level (Fluo4) at point of mPTP opening. **(aii & iii)** Representative traces from the cells treated with 500 nM of WT or A53T respectively. **(aiv)** A53T treated cells open PTP at a lower concentration of Ferutinin than WT (*p* = 0.031). **(bi)** Representative time course images showing that apoptosis (Nucview) is induced followed a dramatic loss of Δψ_m_ after Ferutinin induced PTP opening. **(bii & iii)** Representative traces from WT or A53T treated cells (biv) A53T treated cells induce PTP opening earlier than WT (*p* < 0.0001). **(c & d)** mPTP opening in isolated mitochondria from permeabilized cells. **(ci)** Representative time course images of mPTP opening after applying A53T α-Syn-AF488. **(cii & iii)** Mitochondrial area (ROI 1 area) shows a rapid loss of Δψ_m_ whilst extra-mitochondrial area (ROI 2) shows increased intensity of TMRM after mPTP opens. **(di & ii)** Mitochondria loaded with Fluo-4 and Rhod-5N, after application of A53T and WT. Representative traces and quantitative histogram showing PTP opening occurs earlier in A53T than WT treated mitochondria. *Note.* Data are represented as mean ± SEM. n = 3 – 8 number of wells. Total number of cells = 18 – 80. All experiments were independently repeated 2 – 3 times. *p<0.05, **p<0.001, ***p<0.0001.

We examined whether A53T impairment of respiratory chain function also affects production of ATP, using a genetically encoded mitochondrial FRET-ATP sensor and measured the ratio between yellow fluorescence protein (YFP) and cyan fluorescence protein (CFP) after application of each sample (Figure S5bi). A53T treated cells exhibit significantly reduced ATP production by 42 % compared to WT (Figure 4ei – iii).

Impaired respiration and Complex I function can result in mitochondrial and extra-mitochondrial sources of reactive oxygen species (ROS). Generation of ROS in the cytosol and in the mitochondrial matrix was measured using dihydroethidium (HEt) or MitoCM-H_2_Xros. HEt allows the rate of cytosolic superoxide generation to be measured, as the slope of the ratio of the oxidized form of the dye over the reduced form. Interestingly, application of 500 nM monomeric form of A53T mutant induced an increased rate of superoxide ROS in contrast to WT (WT: 126 ± 5.2 %, A53T: 368 ± 9.1 %, Figure. 4fi – ii). We further showed that the generation of cytosolic ROS is dependent on the concentration of applied monomers (Figure S5c). We then investigated the source of the A53T induced ROS using a range of inhibitors. We found that Mito-TEMPO or MitoQ (mitochondria targeted antioxidants), Trolox (a water - soluble analogue of vitamin E), and Diphenyleneiodonium chloride (DPI) or 4-(2-Aminoethyl)-benzolsulfonylfluorid-hydrochloride (AEBSF) (inhibitors of cytosolic NADPH oxidase (NOX)), effectively blocked A53T induced ROS, suggesting that mitochondrial ROS may be one of the main sources of the excess ROS (Figure 4fiii) with possible activation of NADPH oxidase. MitoCM-H_2_Xros is a reduced non-fluorescent version of Mitotracker red that is fluorescent upon oxidation within mitochondria and therefore the rate also indicates the production of mitochondrial ROS (mROS). We confirmed that A53T induced high levels of mROS (176.6 ± 9.3 % of basal) compared to WT (113.8 ± 4.4 % of basal) (Figure 4gi – iii).

Due to the multiple cellular locations of α-Syn, and the known interaction between α-Syn and lysosomes, we tracked how exogenously applied A53T simultaneously affects mitochondria and lysosomes over time. Following application, A53T α-Syn first induced depolarization and fragmentation of mitochondria, shown by 50 % reduction in TMRM fluorescent signal, while the lysosomal tracker did not change significantly in this time period (Figure S5di – iii), suggesting that the mitochondrial effects of A53T may precede the lysosomal response to A53T.

### A53T induces mPTP opening

We previously showed that early opening of mitochondrial permeability transition pore (mPTP) is a mechanism of α-Syn oligomer induced cell toxicity [12]. We tested whether A53T α-Syn also affects mPTP opening in whole cell and permeabilized mitochondria preparations. Whole cells were loaded with TMRM and the cytosolic calcium dye Fluo4, followed by stepwise application of the electrogenic calcium ionophore Ferutinin, known to induce mPTP opening by mitochondrial calcium overload [29] (reviewed here [30, 31]). mPTP opening was determined by the timepoint of rapid loss of mitochondrial TMRM fluorescence. As demonstrated in Figure 5aii – iv PTP opening occurs at a lower Ferutinin concentration in A53T treated cells (7.6 ± 1.0 μM) than WT treated cells (13.7 ± 1.5 μM). We examined the relationship between mPTP opening and cell toxicity by co-loading TMRM and Nucview488 (an apoptotic marker of caspase-3 cleavage) and applying a high concentration Ferutinin (30 µM) to open mPTP. The large rise in mitochondrial calcium increases mitochondrial permeability and opens the mPTP, and this is then followed by an increase in nuclear green fluorescence demonstrating caspase 3 cleavage and caspase 3 dependent apoptosis. We measured the latency to the rapid loss of TMRM fluorescence after the application of the high concentration of Ferutinin, and observed that application of A53T (74 ± 10 sec) led to early mPTP opening than WT (191 ± 15 sec) (Figure 5bii – iii). Caspase 3 dependent apoptosis was induced in all cells that exhibited mPTP opening.

Next we measured mPTP opening in isolated mitochondria of permeabilized cells. Cells were loaded with Rhod-5N, and permeabilized with 40 µM Digitonin in pseudo-intracellular solution [32]. Application of A53T induced a rapid loss of mitochondrial calcium accumulated dye (Rhod-5N) as the PTP opens and the dye leaves the mitochondria (ROI 1, Figure 5ci & ii), which results in increased Rhod-5N fluorescence in cytoplasm (extra mitochondrial region, ROI 2, Figure 5ci & iii). The number of mitochondria exhibiting mPTP opening within 15 min after the application of α-Syn was higher for A53T compared to WT (Figure 5Dii) in line with the results from the whole cell model.

mPTP opening requires the structural conformation of α-Syn to be a Type-B (beta sheet rich oligomer) [12]. Therefore A53T monomer, in contact with mitochondrial membrane cardiolipin, rapidly forms oligomeric species that are able to open the mPTP, and induce cell toxicity.

### Mitochondrial ROS accelerates oligomerization and A53T induced cell death

We have previously shown that oligomeric α-Syn can induce oxidation of proteins and lipids, both of which can alter PTP opening. We investigated the effect of mitochondrially generated ROS, on the aggregation kinetics and toxicity of A53T within neurons. We measured the oligomer signal (FRET) after co-treating with Trolox, a water-soluble analogue of vitamin E, and Mito-TEMPO, a mitochondria targeted antioxidant. As seen in Figure 6A, application of 100 μM Trolox reduces the intracellular FRET intensity by 30 % (Figure 6aii) but there is no effect on FRET efficiency (Figure 6aiii). The reduction in FRET intensity is due to the reduction of uptake of the A53T monomer across the plasma membrane into cells (Figure 6aiv). However the unaltered FRET efficiency suggests that, while fewer aggregates are formed, those that do form are similar in structure to untreated cells. In contrast, the application of the mitochondrial targeted antioxidant Mito-TEMPO results in significant reduction in both FRET intensity (Figure 6aii) and efficiency (Figure 6aiii) by 27 % and 18 % respectively, with preservation of the donor intensity (i.e. preservation of uptake into cells, but reduced oligomerization). Thus, the generation of mROS is a key factor in the intracellular oligomerization of the A53T α-Syn, and scavenging mROS modulates the formation of aggregates. Finally, we examined the effect of the antioxidants on neuronal cell death shown. Unlike WT (2.9 ± 1.3 %), A53T treated cells induced increased cell death (18.0 ± 3.3 %, Figure 6bi & ii) after 48 hours incubation. A53T induced cell death was effectively inhibited by Mito-TEMPO (17 % reduction to untreated cells, Figure 6ci & ii).

**Figure 6.**
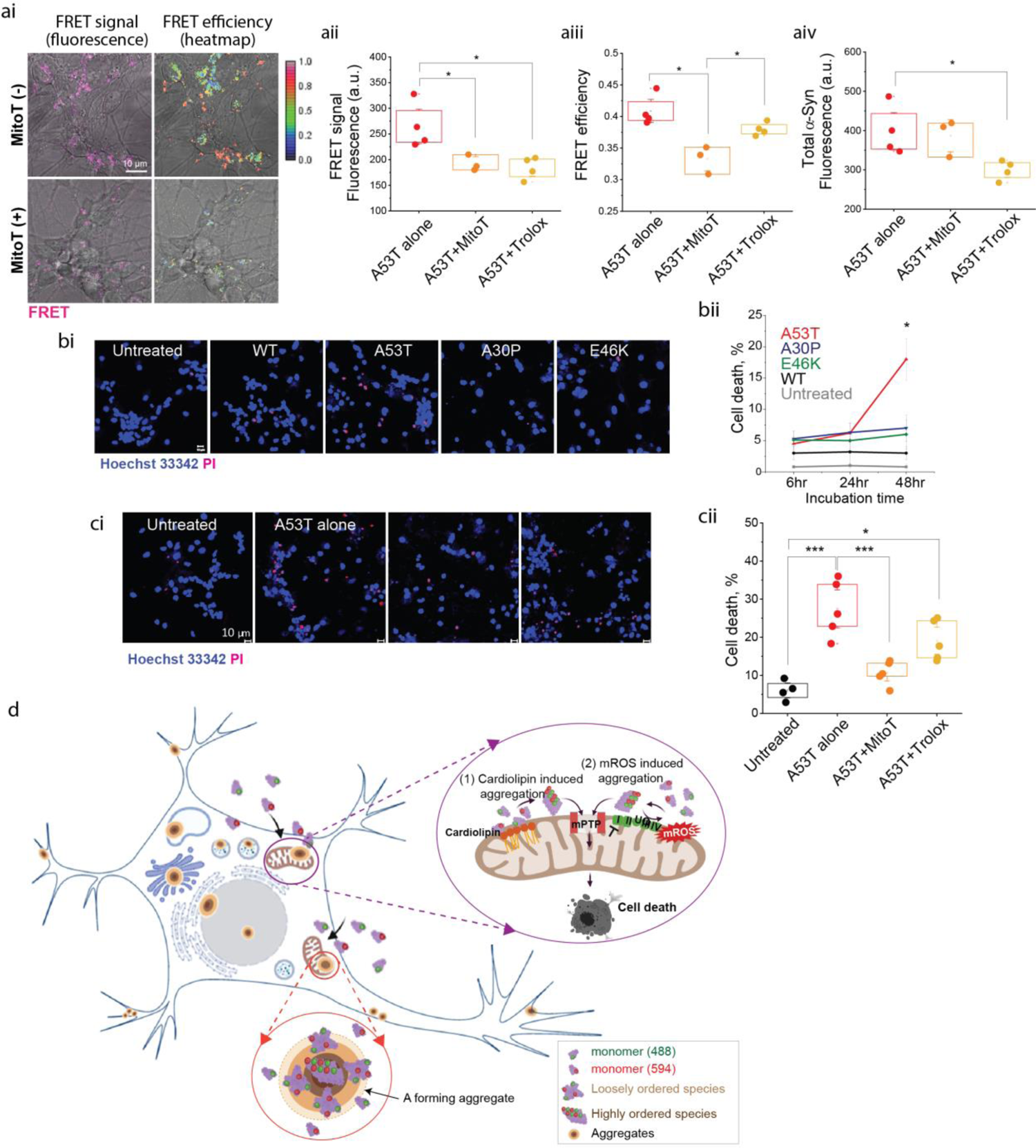
Mitochondrial ROS accelerates oligomerization and A53T induced cell death. **(ai)** Representative images showing both FRET intensity and FRET efficiency of A53T are reduced by treatment with Mito-TEMPO. **(aii & iii)** Mito-TEMPO treated cells show reduced A53T FRET intensity (A53T alone vs Mito-T (Mito-TEMPO): *p* = 0.024, A53T alone vs Trolox: *p* = 0.021) and efficiency (A53T alone vs Mito-T: *p* = 0.011, A53T alone vs Trolox: *p* = 0.039). **(aiv)** Application of Trolox to cells reduces FRET intensity signal through reducing uptake of donor (A53T alone vs Trolox: *p* = 0.020). **(bi & ii)** Cell death induced by 48-hour incubation of A53T (*p* = 0.027), but by WT nor A30P/E46K. **(ci & ii)** A53T induced cell death was rescued by treatment either with Trolox (A53T alone vs Trolox: *p* = 0.018) or Mito-TEMPO (A53T alone vs Mito-T: p < 0.0001). **(d)** Graphical illustration demonstrating how α-Syn monomers form aggregates inside neurons and induce cell toxicity; The intracellular monomeric population begins to self-assemble first into a population of amorphous loosely ordered oligomeric species, and progresses to form highly ordered oligomeric species. Aggregates form with a dense central core of highly ordered oligomer surrounded by a rim of loosely packed oligomers, and occur in multiple hotspots throughout the cell body including the nucleus, Golgi and mitochondria. Mitochondria are a crucial site of aggregation: cardiolipin triggers oligomerization of A53T α-Syn. A53T α-Syn induces over-production of mROS promoting oligomerization of α-Syn. A53T α-Syn oligomerization promotes early opening of mPTP leading to cell death. *Note*. 100 μM Trolox, 0.5 μM Mito-TEMPO were pre-treated 30 min prior to α-Syn application. Data are presented as mean ± SEM. n = 3 – 5 number of wells. Total number of cells = 180 – 491. All experiments were independently repeated 2 – 3 times. *p<0.05, **p<0.001, ***p<0.0001.

These results, therefore, support that mitochondrial dysfunction, in particular with the impaired respiration and generation of mROS, acts synergistically with the effects of cardiolipin, to trigger the intracellular oligomerization of the A53T monomer, and contribute to neuronal death.

### SNCA-A53T hiPSC derived neurons exhibit accelerated seeding and mitochondrial dysfunction

The SNCA-A53T mutation results in an early onset and aggressive disease course caused by a synucleinopathy in cortical and midbrain neurons. We generated human cortical neurons derived from hiPSC from two patients carrying the A53T mutation (see patient details in Figure S6A & Bi & ii) using a modified protocol from [33]. Both control and SNCA-A53T hiPSC derived neuronal cultures are highly enriched (control: 84.3 ± 3.5 %, A53T: 91.3 ± 0.8 % βTubulin positive cells, Figure 7ai) and respond to 5 µM glutamate (control: 71.2 ± 3.1, A53T: 70.3 ± 4.6) (Figure S6ci – iii). We assessed whether SNCA-A53T iPSC derived neurons exhibit aggregate formation, using an ultrasensitive assay that measures the ability for an aggregate to permeabilize membranes and induce calcium influx. The SNCA-A53T iPSC derived neurons both synthesize and secrete higher numbers of oligomeric species than control (Figure 7bi & ii) [34, 35].

**Figure 7.**
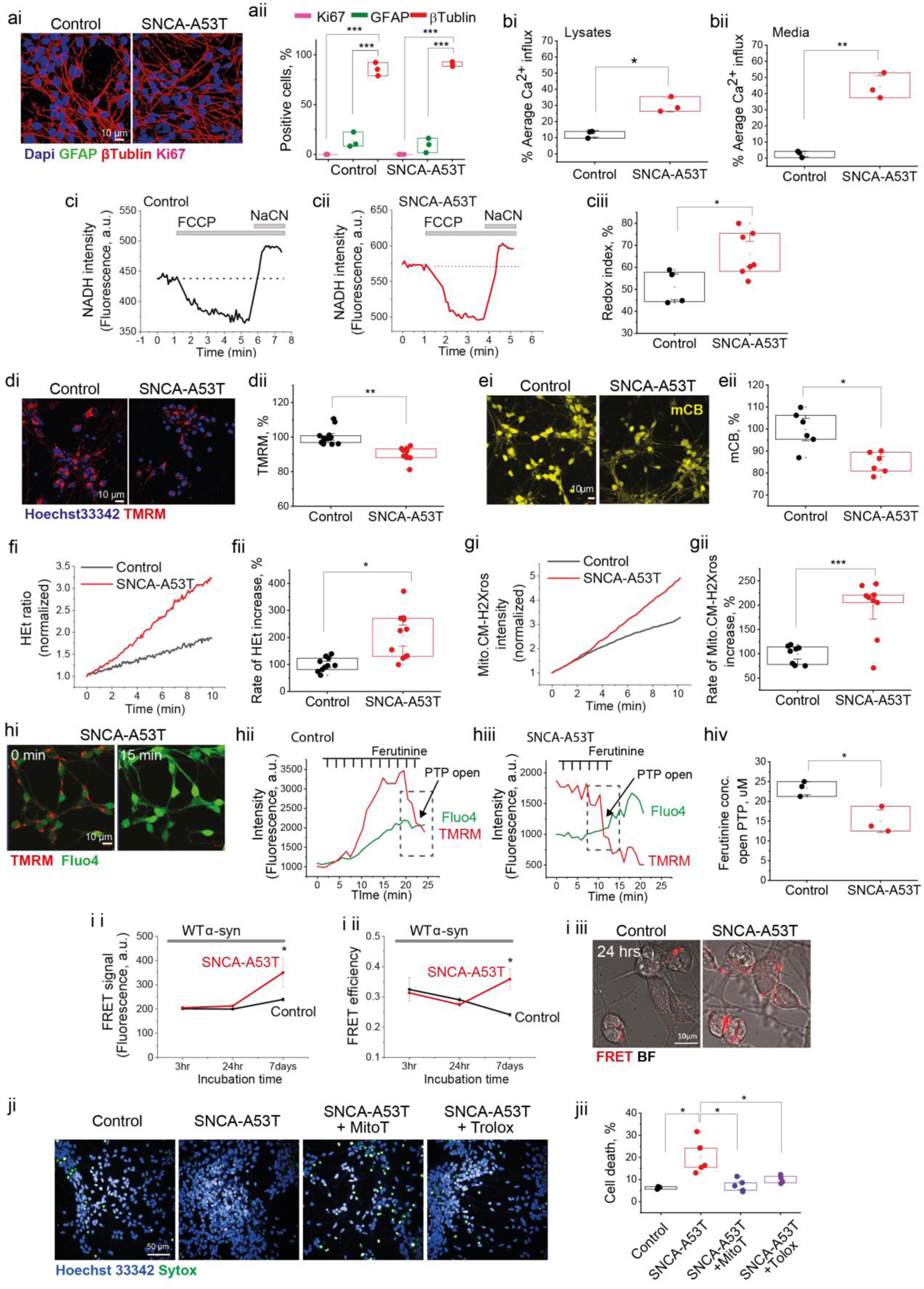
SNCA-A53T hiPSC derived neurons exhibit accelerated seeding and mitochondrial dysfunction. **(ai & ii)** Enrichment of hiPSC derived cortical neurons at day 80, expressing neuronal marker, β tubulin III. **(bi & ii)** Lysates and media from SNCA-A53T neurons contain oligomers that cause increased membrane permeability compared to control neurons. **(ci – iii)** SNCA-A53T neurons exhibit a higher redox index than control neurons, indicating Complex I inhibition (*p* = 0.031). **(di & ii)** There is lower Δψ_m_ in SNCA-A53T neurons than control (*p* < 0.0001), measured by TMRM fluorescence. **(ei & ii)** Glutathione level is lower in SNCA-A53T cells than control (*p* = 0.015). **(fi & ii)** SNCA-A53T neurons exhibit higher ROS production than control (*p* = 0.024). **(gi & ii)** Increased production of mROS measured by MitoTrackerCM-H_2_Xros in SNCA-A53T neurons (*p* < 0.0001). **(hi – iv)** SNCA-A53T neurons open mPTP at lower concentration of Ferutinin (*p* = 0.017). **(i i – i ii)** Application of WT α-Syn-AF488 and α-Syn-AF594 to cells results in higher intracellular FRET intensity and higher FRET efficiency in SNCA - A53T neurons (*p* = 0.039). **(ji & ii)** SNCA-A53T neurons exhibit higher cell death than control neurons, which can be rescued by 0.1 mM Mito-TEMPO and 100 mM Trolox (SNCA-A53T untreated vs Mito-TEMPO (Mito-T): *p* = 0.031, SNCA-A53T untreated vs Trolox: *p* = 0.041). *Note*. Data are represented as mean ± SEM; n = 4 – 8 wells. N = 2 – 3 number of independent inductions. Total number of cells = 321 – 761. All experiments were independently repeated 2 – 3 times. *p<0.05, **p<0.001, ***p<0.0001. See also Figure S6.

Measuring NADH autofluorescence, SNCA-A53T neurons show a significantly higher redox index than control cells indicating that Complex I driven respiration is inhibited in SNCA-A53T cells compared to control (control: 48.0 ± 3.8 %, SNCA - A53T: 33.2 ± 3.8 %, Figure 7Ci – iii). SNCA-A53T neurons also show a reduction in Δψ_m_ (control: 100 ± 1.2 %, SNCA-A53T: 90.1 ± 1.2 %, Figure 7di & ii). Increased cellular oxidative stress was apparent by a reduction in the antioxidant glutathione levels (16.5 % reduction compared to control, Figure 7ei & ii) and an increase in the generation of cytosolic superoxide (206.7 ± 26.9 % increase compared to control, Figure 7fi & ii) as well as mitochondrial ROS (197.3 ± 17.2 % more production compared to control, Figure 7Gi & ii). SNCA-A53T neurons open mPTP at lower concentrations of Ferutinin than control cells (control: 23.3 ± 1.1 µM, SNCA-A53T: 15.0 ± 1.9 µM, Figure 7Hii – iv).

We evaluated intracellular oligomerization by measuring the FRET signal after adding the WT AF488-α-Syn and AF594-α-Syn monomer to SNCA-A53T neurons. SNCA-A53T neurons showed both higher intracellular FRET intensity (control: 239.5 ± 6.9 a.u., A53T: 350.1 ± 60.4 a.u. at 7 days, Figure 7ii) and efficiency (control: 0.24 ± 0.007, SNCA-A53T: 0.35 ± 0.03 at 7 days, Figure 7Iii) compared to control cells. We also measured FRET after adding AF488/AF594 oligomers, and found that the addition of WT oligomers causes a rapid increase in both FRET intensity and efficiency, demonstrating that the SNCA-A53T cells are able to effectively and rapidly seed aggregation. (Figure S6j). Finally, Figure 7Ji & ii shows that the mutant cells exhibit a higher level of basal cell death (control: 6.9 ± 0.3 %, SNCA-A53T: 20.1 ± 3.4 %), that is prevented with both 100 µM Trolox (10.0 ± 1.0 %) and 0.1 µM Mito-TEMPO (7.3 ± 1.6 %) after 24-hour incubation.

Taken together, cortical neurons derived from PD patients expressing the A53T mutation display bioenergetics defects, and oxidative stress, and this results in accelerated seeding, oligomerization, PTP opening and cell death.

## Discussion

Precise characterization of the dynamic process of protein aggregation within human cells is a major challenge, leaving the causal mechanisms for the initial stages of aggregation in disease largely unknown. Characterizing which factors favour or hinder aggregation, and how toxic aggregate species interact within neurons to drive neuronal cell death is fundamental to understanding PD pathogenesis. Here we describe the kinetics of intraneuronal aggregation of α-Syn, how and where structural conversion occurs in the human neuron, and the functional consequences of this conversion.

Our understanding of the kinetics of α-Syn aggregation has emerged largely from *in vitro* buffer experiments [10, 16, 17, 20, 36]. Aggregation commences with a lag phase due to the difficulty of monomers alone assembling to form oligomers (primary nucleation), followed by a dramatic growth phase in which conversion is accelerated, due to the relative ease of adding monomers to existing seeds, or oligomers, to form protofilaments and fibrils (elongation). Such kinetic models however do not consider the complexity of the cellular environment, with the presence of cellular organelles, lipid surfaces, and altered pH, and protein degradation systems. smFRET offers a powerful, non-invasive approach to investigate individual molecular species and aggregation and has been used successfully to study protein interaction and conformation in live cells [15, 37]. We applied smFRET to directly detect and characterize the kinetics of formation, interconversion and accumulation of externally applied oligomers from its early monomeric state, in the cellular environment. smFRET was able to provide estimations of the size (deduced from fluorescence intensities) and structural group (deduced from FRET efficiencies) for each individual oligomeric species detected [10, 38]. Inside human neurons, the monomeric population begins to rapidly self-assemble. We observed two distinct populations-amorphous oligomers, termed Type-A oligomers (characterized by low FRET efficiency), were loosely packed, and the first to form initially, and, these progress to form Type-B oligomers (characterized by higher FRET efficiency), which are more compact, have a higher order β-sheet structure over days. The kinetics of self-assembly and oligomer formation were dependent on the initial concentration of monomer, and time inside the cells, and the sequence of the protein, in keeping with the known critical factors that affect aggregation *in vitro*.

The A53T mutation alters the kinetics of intracellular aggregation, with abolition of the lag phase, resulting in immediate self-assembly into small oligomers, and rapid accumulation of more Type B oligomers, in line with *in vitro* data [9, 16]. The dramatically altered aggregation kinetic profile in cells is likely due to the abundant protein-lipid interactions inside the cell. α-Syn is disordered in solution but can adopt an α-helical conformation upon binding to lipid membranes. The interaction of α-Syn with lipids at membrane surfaces can trigger conversion of α-Syn from its soluble state into an aggregated state, enhance the rate of nucleation, and significantly reduce the lag phase [39]. Lipid-induced generation of fibrils is highly sensitive to the specific sequence of the protein, in particular, the region encompassing the residues 46 – 51, and the rate of lipid-induced aggregation is significantly higher in A53T compared to WT protein [39]. Our data suggests that human neurons, rich in multiple membrane surfaces, including mitochondrial membranes, can trigger the aggregation of A53T by stimulating primary nucleation, and that the lipid-protein interaction is heavily dependent on the sequence of the protein, with A53T enhancing the lipid interaction.

We used 3D FRET-CLEM to precisely characterize the morphology and location of the different A53T species inside cells. CLEM is an integrated approach, that closes the spatial and temporal resolution gap between light microscopy observations and EM, by enabling the overlay of fluorescence and electron microscopy images from the same cell. This allowed us to complement dynamic information from live cell imaging with high-resolution ultrastructural information, providing unique data on the location of a fluorescently labelled molecule within the ultrastructural context of a cell (Anderson et al., 2019). Due to the challenge of imaging multiple landmark fluorophores in addition to FRET (related to phototoxicity for human neurons), overlay was performed using the nucleus alone. The accuracy of the overlay is limited because the nucleus is a large structure compared to the target FRET oligomer, and, given the shrinkage and warping that occurs during sample preparation for EM, this can result in high uncertainty in the overlay. Notwithstanding these limitations, we observed first, a stereotypical evolution of an aggregate forming with a dense central core of high FRET intensity, surrounded by a rims of lower FRET intensity, throughout the neuronal soma and processes. Second, aggregation clearly has a heterogeneous location, occurring in multiple hotspots throughout the cell body, but often in crowded cellular environments rich in organelles, including membrane surfaces from the nucleus, plasma membrane, Golgi, and mitochondria. Thus the dramatic increase in aggregation kinetics of α-Syn inside the cell, is related to multifocal seeding events occurring at different membrane surfaces, accelerating primary and secondary nucleation processes. This provides a possible explanation of the way that the key initial steps in the process of aggregation of α-Syn could occur in a cellular environment and ultimately lead to the onset and proliferation of disease affecting specific organelles.

Mitochondrial dysfunction is a key player involved in the pathogenesis of sporadic and familial PD [40]), and an interaction between mitochondria and synucleinopathy has been reported [23, 41, 42]. α-Syn has been shown to selectively bind to mitochondrial membranes (Chinta et al., 2010; Devi et al., 2008; Guardia-Laguarta et al., 2014; W. W. Li et al., 2007; Liu et al., 2009; Ludtmann et al., 2018; Shavali et al., 2008; Wang et al., 2019), and may play a physiological role in mitochondrial function, mediating maintenance of mitochondrial fission, Complex I activity and calcium signaling (reviewed here [43, 44]). In contrast to previous studies largely demonstrating mitochondria as an effector of α-Syn oligomers, here we demonstrate two key mechanisms by which the mitochondria may be a key organelle that seeds the aggregation of α-Syn. Mitochondrial membranes are made up of a range of different phospholipids, although cardiolipins are specific phospholipids of the mitochondria, comprising about 20 % of the inner mitochondria membrane phospholipids mass, where it maintains the structural and functional integrity of proteins and enzymes involved in the mitochondrial electron transfer chain and oxidative phosphorylation [45]. Translocation of cardiolipin to the outer mitochondrial membrane has been reported, and externalized cardiolipin is involved in the modulation of mitophagy. Interestingly α-Syn has been demonstrated to interact with cardiolipin, and mutations also induce the externalization of cardiolipin to the outer mitochondrial membrane [46].

In our study, we show that (i) α-Syn binds cardiolipin and adopts an alpha-helical conformation when bound to cardiolipin (ii) cardiolipin is a potent trigger for the rapid aggregation of A53T α-Syn to form fibrils, without a lag phase *in vitro* (iii) during cardiolipin induced aggregation, the cardiolipin is sequestered within the protein aggregate as it forms, and this lipid-protein co-assembly acts as a reactant for further surface-induced elongation. Previous studies using NMR and cryo-EM highlight how lipids co-assemble with α-Syn molecules, implying strong lipid-protein interactions [47]. End stage aggregates are now known to be composed of a multitude of organelles, lipid membranes and mitochondria [48]. Therefore lipids both trigger aggregation, and act as reactant during aggregation. Our data would support a mechanism by which the aggregate starts at a (cardiolipin containing) membrane surface and sequester the lipid within it as the aggregate grows into a mature structure.

In addition to providing the lipid seed for aggregation, mitochondria are a critical source of reactive oxygen species (mROS) generated through mitochondrial metabolism and the electron transport chain. Impairment of mitochondrial function can lead to changes in the redox state and regulation of the cell. Here we show that A53T α-Syn inhibits Complex I dependent respiration, impairs ATP production, and depolarizes the mitochondrial membrane. This bioenergetic deficit leads to an increase in both mitochondrial and cytosolic generation of ROS. The oxidative environment of dysfunctional mitochondria in turn promotes further oligomerization of A53T monomers in neurons. Inhibition of mROS formation, using mitochondrial free radical scavengers, can suppress A53T oligomerization and abolish A53T induced toxicity. We have previously demonstrated how oligomers, once formed, interact with metal ions to generate further ROS, and ultimately oxidative stress [49]. Lipid peroxidation, particularly of polyunsaturated fatty acids (PUFAs), from aberrant ROS formation is a major inducer of α-Syn oligomer toxicity through enhancing the lipid-aggregate interaction [34, 50]. Cardiolipin contains a high number of PUFAs, which makes it an easy target of peroxidation. Abnormal cardiolipin content, fatty acyl chain composition, and level of oxidation are associated with a significant decrease of membrane potential and multiple defects in mitochondrial function (reviewed here [51]). Taken together, our results show that inside cells, the effect of mROS is bi-directional, and self-amplifying: mROS promotes oligomerization of α-Syn, and oligomer formation impairs Complex I function and induces further mROS generation. In summary, whilst the physical interaction of A53T and cardiolipin is sufficient to trigger aggregation, the presence of mROS within mitochondria can drive the aggregation reaction.

The final common pathway of A53T within the mitochondria is the altered permeability of the mitochondrial outer membrane. We previously reported that WT α-Syn oligomers of the Type -B structure, are located at the mitochondrial membrane in close proximity to ATP synthase and Complex I. At this site, oligomeric species of α-Syn can generate the production of free radicals, that induce local oxidation events in mitochondrial lipids, resulting in mitochondrial lipid peroxidation, and mitochondrial proteins, and together, these events lead to mitochondrial membrane permeabilization [12]. In this study, we demonstrate that the application of A53T monomer to isolated mitochondria or whole cells, induces rapid permeability transition of the mitochondrial membrane, similar to the effect of the WT α-Syn Type-B oligomer. This data, taken together with the cardiolipin induced aggregation, raises the hypothesis that A53T contacts cardiolipin, triggering oligomer formation and conversion to Type-B oligomers, resulting in oligomer induced impaired respiratory chain function and reduced ATP production, oligomer induced mROS production, and oligomer induced mitochondrial membrane depolarization. Together these effects predispose mitochondria to early opening of the permeability transition pore and activation of caspase 3 dependent apoptosis.

We utilized hiPSC-derived neurons bearing SNCA-A53T mutation to model the effect of the aggregation of endogenous α-Syn in disease. hiPSC derived from patients with A53T mutations have been used successfully to generate neurons that exhibit PD relevant phenotypes including α-Syn accumulation and mitochondrial dysfunction [52–54]. In agreement with our findings that A53T exhibits increased oligomerization, we identified higher levels of oligomers in the SNCA-A53T neurons. Increased aggregation in SNCA-A53T was associated with impaired respiration, depolarized mitochondrial membrane potential, and oxidative stress. The A53T cellular environment, with intracellular aggregates and oxidative stress together resulted in accelerated oligomerization of exogenously added α-Syn. Finally, the presence of A53T oligomers was sufficient to induce early opening of the mPTP, and result in cell death. Removal of one of the major drivers of mPTP and oligomerization, mROS, was able to prevent the cellular pathology in the SNCA-A53T model, thereby lending support for our hypothesis that mitochondrial ROS is a key driver in α-Syn pathology.

## Conclusion

Our study has integrated high resolution biophysical approaches with human iPSC biology to comprehensively characterize the spatial and temporal features of protein aggregation in the intracellular environment. We were able to show that oligomerization of the A53T α-Syn occurs rapidly in the human neuronal environment, and preferentially on membrane bound structures. Oligomerization can occur on mitochondrial membranes, where it is triggered by cardiolipin and promoted by mROS. Mitochondrial oligomerization causes pathological effects on the bioenergetics state of the cell, evidenced by reduced Complex I activity and increased oxidative stress, ultimately leading permeabilization of mitochondrial membranes, mitochondrial dysfunction and cell death. Thus, we highlight the crucial role of mitochondria in underpinning A53T aggregation and the toxicity, indicating that a synergistic interaction between α-Syn and mitochondrial membranes may be the common downstream pathological mechanism in synucleinopathies.

## Acknowledgements

This work was supported by the Wellcome/MRC Parkinson’s Disease Consortium grant (grant number WT089698) and the National Institute of Health Research University College London Hospitals Biomedical Research Centre. SG was supported by Wellcome, and is an MRC Senior Clinical Fellow (MR/T008199/1). This research was funded in part by Aligning Science Across Parkinson’s [Grant number: ASAP-000509] through the Michael J. Fox Foundation for Parkinson’s Research (MJFF). The single-molecule instruments used in this study were supported by the UK Dementia Research Institute, UCB Biopharmaceuticals, Alzheimer’s Research UK (ARUK-EG2018B-004), and a kind donation from Dr. Jim Love. MH and AC were supported by the Engineering and Physical Sciences Research Council (EPSRC) and Medical Research Council (MRC) through the CDT in Optical Medical Imaging (OPTIMA), grant number EP/L016559/1, and the Rosetrees Trust. LT is supported by Kennedy’s Disease Association and by AFM-Telethon.

## Author contributions

Conceptualization, M.L.C. and S.G.; Methodology, S.G., M.H.H., L.C., and A.Y.A.; Investigation, M.L.C., A.C., B.P.S., C.M., A.V.B., S.D., C.P., D.A., G.S.V., W.Z. and J.R.E., A.I.W., Z.H.Z., A.T.; Resources, M.R., E.F., P.R.A., N.E. K.M., L.T., D.L., P.G., D.K. and T.K.; Equally contributed second authors, A.C. and B.P.S.; Writing-original Draft, M.L.C.; Writing-review & Editing, S.G., D.A., A.Y.A and M.H.H; Funding Acquisition, S.G.

## Methods

### Human recombinant α-Syn

#### Aggregation of human recombinant α-Syn

Monomeric WT α-Syn was purified from *Escherichia coli* as previously described in [55]. Aggregation reactions were carried out using a solution of α-Syn 70 μM in 25 mM Tris buffer supplemented with 100 mM NaCl, pH 7.4 (in the presence of 0.01 % NaN_3_ to prevent bacterial growth). The buffer was freshly prepared before each experiment and passed through a 0.02 μm syringe filter (Anotop, Whatman) to remove insoluble contaminants. Prior to incubation, the reaction mixture was ultra-centrifuged at 90k g. for 1h at 4°C to remove potential seeds. The supernatant was collected and separated in two fractions: one kept at 4°C at all times until use (monomers), and a second incubated in the dark at 37°C and 200 r.p.m., during ∼7 – 8 hours to avoid fibril formation (monomers + oligomers). α-Syn was always kept in LoBind microcentrifuge tubes (Eppendorf, Hamburg, Germany) to limit surface adsorption.

#### Labelling of α-Syn WT, A53T, A30P and E46K

The A90C mutant variant of α-Syn WT and mutants (A53T, A30P and E46K) was purified as a monomeric fraction from *Escherichia coli* as described previously [55] and labelled with either maleimide-modified AF488 or AF594 dyes (Invitrogen, Carlsbad, CA, USA) via the cysteine thiol moiety as previously reported [56]. The labelled protein was purified from the excess of free dye by a P10 desalting column with Sephadex G25 matrix (GE Healthcare, Waukesha, WI, USA), divided into aliquots, flash frozen, and stored at −80°C [10]. Each aliquot was thawed immediately and used only once. The reaction yield was checked by mass spectrometry for all reactions, and all labeling reactions with a yield lower than 90% were discarded.

#### Aggregation of labelled α-Syn (WT, A53T, A30P and E46K)

Protocol is similar to the unlabelled version of the protein except that the collecting time point to obtain oligomer is after 30hr due to slower aggregating [10].

### Cell culture

#### Primary rat cortical co-culture

1 – 3 days postpartum Sprague Dawley rats (University College London breeding colony) were used and experimental procedures were performed according to the United Kingdom Animal (Scientific Procedures) Act of 1986. Rat cortices were placed in an ice-cold dissecting buffer (HBSS supplemented with 10 mM HEPES and 20 % FBS) and washed five times with a washing buffer (HBSS supplemented with 10mM HEPES). The tissue was digested with 0.5 % EDTA-trypsin supplemented with DNAse for 15 min and neutralized with the dissecting buffer. After washing twice again with the washing buffer, the tissues were dissociated with washing buffer supplemented with DNAse and then the pellets were collected in Neurobasal completed medium (Neurobasal A medium supplemented with B27, 2 mM Glutamax, Pen/Strep). Approximately 600,000 cells were plated on 25 mm coverslips (PDL coated) and or 200,000 cells for 8-well ibidi chambers (PDL coated). The cultures were maintained at 37 ‘C (5% CO2) and the media were changed every 4 – 5 days. Cells were used at 12-16 days.

#### hiPSC culture

hiPSCs were derived from donors who had given signed informed consent for derivation of hiPSC lines from skin biopsies as part of the EU IMI-funded program StemBANCC and reprogrammed as described [57, 58]. Briefly, Cyto Tune-iPS reprogramming kit (ThermoFisherScientific) was used to reprogram fibroblasts through expression of OCT4, SOX2, KLF4 and c-MYC by four separate Sendai viral vectors. Control 1 and 2 were derived by StemBANCC from an unaffected volunteer and control 3 was purchased from ThermoFisherScientific. hiPSCs were generated from familial Parkinson’s disease patients carrying a point mutation in SNCA (A53T point mutation). Presence of mutations were confirmed by a sanger sequence (GENEWIZ). hiPSCs were maintained on Geltrex in Essential 8 medium (ThermoFisherScientific) and passaged using 0.5 mM EDTA.

#### Differentiation into cortical neurons

Neural induction and differentiation were performed using a modified version from [33]. Briefly, upon 100 % confluence of hiPSC, SB431542 (10 µM, Tocris) and Dorsomorphin (1 µM, Tocris), dual SMAD inhibitors were treated in N2B27 medium (DMEM:F12, insulin, 2-mercaptoethanol, Non Essential Amion Acids, N2 supplement, Pen/Strep, Neurobasal, B27, Glutamax, Pen/Strep) and medium was refreshed every day for 10 – 12 days until neuroepithelium appear. Then the neuroepithelium sheets were split using dispase and cultured in N2B27. At around 35 days of induction, cells were dissociated into a single cell using accutase and approximately 150,000 number of cells plated on either PDL and laminin coated glass bottom 8-well slide chambers (Ibidi/Thistle, cat No.80826), Geltrex coated 8-well ibidi chambers (cat No. IB-80826) or 96-well plates (Falcon, cat No. 353219). Medium was replaced every 4 – 5 days and cells were used at 60 – 90 days after induction.

### Live cell imaging

Live cell imaging was performed using an epi-fluorescence inverted microscope equipped with a CCD camera (Retiga; QImaging) or confocal microscope (Zeiss LSM 710 or 880 with an integrated META detection system). For epi-fluorescence inverted microscope, excitation was provided by a xenon arc lamp with the beam passing through a monochromator (Cairn Research) and emission was reflected through a long-pass filter to a cooled CCD camera and digitized to 12-bit resolution (Digital Pixel Ltd, UK) and the data were analyzed using Andor iQ software (Belfast, UK). For confocal microscopes, illumination intensity was limited to 0.1 – 0.2 % of laser output to prevent phototoxicity and the pinhole was set to allow optical slice at approximately 1 – 2 μm. Pre-room temperature warmed HBSS (supplemented with 10 mM HEPES and adjusted at pH 7.4) was used as a recording buffer. 3 – 6 fields of view per well and at least 3 wells per group were used to analyze using ZEISS ZEN software, Volocity 6.3 cellular imaging or ImageJ software. All experiments were repeated at least 2-3 times with different animal batches (primary co-culture) and inductions (hiPSC derived neurons).

### Measurement of Oxidative stress

*Superoxide*; cells were washed and loaded 2 uM dihydroethidium (HEt, Thermo Fisher Scientific) in the recording buffer. HEt is an indicator of superoxide which exhibits blue fluorescence in the cytosol before oxidation and the nucleus presents a red fluorescence upon oxidation. HEt allows the rate of superoxide generation to be measured which is present as the ratio of the oxidized form of the dye over the reduced form. Recording was performed using an epi-fluorescence inverted microscope equipped with 20x objective after a quick loading (2 – 3 min) in order to limit the intracellular accumulation of oxidized product and the dye was present throughout the imaging. Excitation was set up to 530 nm and emission recorded above 560 nm was assigned to be for the oxidized form while excitation at 380 nm and emission collected from 405 nm to 470 nm were for the reduced form. The ratio of the fluorescence intensity resulting from its oxidized/reduced forms was quantified and the rate of ROS production was determined by dividing the gradient of the HEt ratio after application of recombinant α-Syn against basal gradient.

*Mitochondrial ROS*: it was assessed using MitoTracker® Red CM-H2XRos dye, a reduced non-fluorescent version of Mitotracker red that is fluorescent upon oxidation within mitochondria, (Thermo Fisher Scientific) which accumulates into mitochondria upon oxidation. Cells were washed twice and loaded with 500 nM MitoTracker® Red CM-H2XRos for 20 - 30 min with the dye in the recording buffer at room temperature. The fluorescence measurement was obtained by using a confocal microscope with a 40x oil-immersion objective lens excited with 561 nm laser and emission detected above 580 nm. The rate of increase in red fluorescence for each cell was analyzed as the production of mitochondrial ROS.

### Mitochondrial membrane potential (ΔΨm)

*ΔΨm* is the electrogenic potential between the inner membrane and matrix of mitochondria which combines with the mitochondrial pH gradient provides the force driving protons into mitochondria to generate ATP. Cells were washed twice and loaded with 25 nM tetramethylrhodamine methyl ester (TMRM, Thermo Fisher Scientific), a lipophilic cationic dye that accumulates within mitochondria in inverse proportion to Δψ_m_ according to the Nernst equation, in the recording buffer for 40 min at room temperature. Using confocal microscope imaging was performed in the present of TMRM for which 560 nm laser was used to excite and fluorescence was measured above 580 nm. Z-stack images were collected and the fluorescence intensity of TMRM was analyzed using ZEISS ZEN software (Zeiss). The assessment of how mitochondria maintain the potential was obtained on a single focal plane (time series) by application of 2 μg/ml Oligomycin (inhibitor of Complex V), 5 μM Rotenone (inhibitor of Complex I) and 1 μM FCCP (mitochondrial uncoupler).

### [Ca^2+^]c imaging

This was measured using Fura-2, AM which is a ratiometric dye with a high affinity for Ca^2+^. The cytosolic Ca^2+^ as well as the rapid transient kinetics and decay times were assessed. 5 uM Fura-2 was loaded with 1.25uM Pluronic acid for 40 min and then washed twice before imaging. The fluorescence measurement was obtained on an epifluorescence inverted microscope equipped with a 20x objective. [Ca^2+^]c was monitored in a single cell by obtaining the ratio between the excitation at 340 nm (high Ca^2+^) and 380 nm (low Ca^2+^) for which fluorescence light was reflected through a 515 nm long pass filter.

### NADH redox state and total NADH pool

NADH autofluorescence was measured as described in [27, 59] using an epi-fluorescence inverted microscope equipped with a 40x oil-fluorite objective with an excitation at a wavelength of 360 nm within 455 nm emission. NADH redox state indicates the balance between electron transport chain (ETC) activity and the rate of substrate supply as a key mechanism to maintain functional mitochondria such as how efficiently cells produce ATP. In the cells, the resting level of NADH is expressed in the form of autofluorescence and the NADH redox state can be measured as a function of the maximally oxidized and maximally reduced. Increased oxidation of NADH is associated with decreased NADH autofluorescence, whereas more reduced state shows increased NADH autofluorescence that is associated with impaired respiration.

*Measurement of NADH redox state*; 1 μM FCCP enables to stimulate maximal respiration by completely oxidizing all mitochondrial NADH into NAD+ and H+ to produce maximum ATP which resulted in minimal autofluorescence. Following stabilization of the signal, 1 mM NaCN was added which inhibited the respiration as preventing NADH oxidation, and as a result maximal NADH reduction was obtained representing maximal autofluorescence. The total NADH pool was calculated by subtracting the minimal autofluorescence (by 1 uM FCCP) from the maximal signal (by 1 mM NaCN). The NADH redox state was expressed as a percentage of basal in total NADH. Higher redox state indicates more reduced NADH which can suggest Complex I driven respiration is inhibited [27, 60].

### ATP Measurement

Cells were transfected with the mitochondrial-targeted ATP indicator AT1.03 [61] that allows visualization of the dynamics of ATP. The ratio between yellow fluorescence protein (YFP) and cyan fluorescence protein (CFP) was measured after application of each monomer sample. When ATP is not bound to the probe, low FRET is detected which is emitted from the CFP whilst upon binding ATP to the probe, the two fluorescence proteins are close to generate FRET which results in YFP signal. Cells were excited with 405 nm laser and emission for CFP is at 460 – 510 nm and 540 – 600 nm for YFP.

### Mitochondria permeability transition pore (mPTP) opening

It is performed as described in [32]. Briefly, cells were washed twice and loaded with 25 nM TMRM and either 5 μM Nucview 488 (Biotium) or 5 μM Fluo4-AM (Thermo Fisher Scientific). The fluorescence signals were measured with laser 488 nm for Nucview 488 and Fluo4 and emission between 488 nm and 516 nm and 560 nm laser with emission above 580 nm for TMRM after application high concentration of Ferutinin (30 µM) at once or 2 μM Ferutinin in a stepwise manner that leads to mPTP opening upon reaching the threshold. The threshold of mPTP opening was measured as the time point at which rapid loss of TMRM fluorescence occurred after applying Ferutinin or α-Syn samples. To permeabilize cells, cells were exposed to 40 – 60 μM digitonin in a pseudo-intracellular solution consisting of 135 mM KCL, 10 mM NaCl, 20 mM HEPES, 5 mM pyruvate, 5 mM malate, 0.5 mM KH_2_PO_4_, 1 mM MgCl_2_, 5 mM EGTA, and 1.86 mM CaCl_2_ yielding approximately 100 nM [Ca^2+^].

### Cell death assay

Cell death was detected using Propidium iodide (PI, Thermo Fisher Scientific) or SYTOX™ Green (SYTOX, Thermo Fisher Scientific) which is excluded from viable cells but exhibits red fluorescence following a loss of membrane integrity and Hoechst 33342 (Hoechst, Thermo Fisher Scientific) which stains chromatin blue in all cells to count the total number of cells. 20 μM PI or 500 nM SYTOX and 10uM Hoechst were directly added into the dishes and cells were incubated for 15 min. The fluorescent measurements were using a confocal microscopy. Hoechst and PI were excited by 405 nm and 543 nm laser lines with the emission between 405 nm and 470 nm (Hoechst) and 570 nm to 640nm (PI) respectively. SYTOX was excited by 488 nm laser with the emission between 488 nm and 516 nm. Percent cell death was quantified by the percent between the number of red fluorescent cells in total number of Hoechst 33342 expressing cells per image.

### Intracellular FRET measurement

Intracellular FRET was measured using a confocal microscope. We used AF488 and AF594 as a FRET pair and the equimolar concentration was treated in cells grown in 8-ibidi chambers (cat number IB-80826). After a required period of time, cells were washed twice and replaced with the recording buffer. As a control, either AF488- or AF594-only labelled treated cells were also measured under the same conditions. As a measure of total α-Syn, the AF594-labelled α-Syn was directly excited and its intensity measured (the AF488 was not used for this, since its intensity is also affected by the FRET process). As a measure of aggregated species, AF488 was excited by 488 nm irradiation, emission detected from AF594.

The intracellular FRET efficiency was calculated using a custom-written script in Igor Pro (Wavemetrics). Briefly, following subtraction of the autofluorescence, the images were thresholded using the directly excited AF594 channel. The FRET efficiency for each voxel containing AF594 intensity was then calculated according to Equation 1 [20, 62].

### Single-molecule confocal microscopy

Cells were treated with AF488 and AF594 labelled monomers and incubated for various time points. Then the lysates were collected using a lysis buffer (150 mM sodium chloride, 1 % Triton-X, 50 mM Tris pH 8.0) and analyzed using single-molecule confocal microscopy [20].

The samples were first diluted to concentrations ∼50 pM before being loaded into a 200 μL gel-loading tip (Life Technologies, Carlsbad, CA, USA) attached to the inlet port of a microfluidic channel (25 μm in height, 100 μm in width, 1 cm in length) mounted onto the single-molecule confocal microscope (described below). The confocal volume was focused 10 μm into the center of the channel, and the solution was passed through the channel at an average velocity of 2 cm/s by applying a negative pressure, which was generated using a syringe pump attached to the outlet port via Fine Bore Polyethylene Tubing (0.38 mm inner-diameter, 1.09 mm outerdiameter; Smiths Medical International, Hythe, Kent, UK). After the appearance of single-molecule bursts corresponding to labelled α-Syn passing through the confocal volume, the sample was measured for 600s.

The confocal microscope is similar to those used previously (Horrocks et al., 2015). A Gaussian laser beam at 488 nm (100 mW, LBX-488-100-CSB-OE, Oxxius) was directed through the back-port of an inverted microscope (Nikon Te2000-U). A dichroic mirror (DI03-R405/488/561/635, Semrock) reflected the beam through an oil-immersion objective lens (Nikon CFI Plan Apochromat VC 100x Oil, NA 1.4, W.D 0.13 mm) which focused to a diffraction-limited confocal spot. The emitted fluorescence was collected by the objective lens and was passed through the same dichroic before being focused by the tube lens through a 50 µm pinhole (Thorlabs). The AF488 and AF594 fluorescence were separated by a second doichroic filter (LP02-647RU-25, Semrock). The AF594 fluorescence was passed through a band-pass filter (FF01-629/53, Semrock) before being focused onto an Avalanche Photodiode Detector (APD) (PerkinElmer). The AF488 fluorescence passed through a second filter set (long-pass: BLP01-488R-25 and band-pass: FF01-525/30-25, Semrock) onto a second APD. Outputs from the APDs were connected to a USB data acquisition card (USB-CTR04, Measurement Computing), which counts the signals and combines them into time-bins of 50 μs, the expected residence time of the molecules in the confocal volume. The data were analyzed as in [20] using custom-written scripts.

### Total internal reflection fluorescence microscopy (TIRFM)

The dual labelled α-Syn samples were diluted to nanomolar concentrations to facilitate single-molecule imaging. Borosilicate glass coverslips (20 × 20 mm, VWR International, USA. Product number 63 1-0122) were cleaned using an argon plasma cleaner (Zepto, Diener, Germany) for 40 minutes to remove any fluorescent residues. Frame-Seal slide chambers (9 × 9 mm^2^, Biorad, Hercules, USA. Product number SLF-0601) were affixed to the glass, and the samples were added to the cover slide on the inside of the chamber and incubated for at least 30 minutes, before being washed with 20 nm filtered buffer (20 mM Tris, 100 mM NaCl, pH 7.1). Imaging was performed using a home-built, bespoke single-molecule total internal reflection fluorescence (TIRF) microscope described previously [63, 64].

Fluorophore-labelled samples were illuminated with 488 nm laser (∼50 W cm^-2^) and the intensity of the emission from AF488 measured over a time period of 10 seconds and recorded as a TIFF stack. Only those species that also had intensity resulting from AF594 due to FRET were analyzed. All image processing was done using ImageJ. For each TIFF stack, the species were located in the first slice using ImageJ’s find maxima function using a prominence level selected to identify all fluorescent puncta. The intensity of each of the located maxima was extracted for each slice in the TIFF file. These data were then analyzed using a custom-written script (MATLAB). Chung-Kennedy filtration was performed on the resulting intensity traces using a window size of 8. An approximate differential of the filtered data was calculated, and the peaks associated with the discontinuities at each step located using a threshold of 455 nm. The validity of the located steps was assessed by calculating the t-statistic, the ratio of individual step height to the local regional variance, and ensuring that this was above a threshold level of 0.01. Traces in which it was not possible to discern individual steps were removed from analyses. Since only the number of AF488 fluorophores were quantified, the size of the oligomer was determined by multiplying the total number of steps by two.

### Correlative Light and Transmission Electron Microscopy (CLEM)

Cells were plated and grown in 35 mm gridded glass-bottom dishes (P35G-1.5-14-CGRD, Mattek). At the appropriate timepoint, an equal volume of fixative (8 % Formaldehyde in 0.2 M phosphate buffer (PB), pH 7.4), pre-warmed to 37°C, was added directly to the growth medium and left for 15 mins. Cells were then washed in 0.1 M PB, and confocal images using Zeiss LSM 880 (x25, x65 objective lenses) then acquired from the area of interest that is identified based on a single cell with healthy morphology. Samples were then post-fixed in 2.5 % Glutaraldehyde/ 4 % Formaldehyde in 0.1 M PB (pH 7.4) for 30 mins, stained in 1 % reduced osmium (1 % osmium/1.5% potassium ferricyanide) at 4 °C for one hour, followed by 1 % tannic acid for 45 mins at room temperature (RT), prior to quenching in 1% sodium sulphate at RT for 5 mins and washing in double distilled water (3 x 5 mins) at RT. The glass coverslip was then removed from the Mattek dish, and dehydrated through a graded series of ethanol (25 %, 50 %, 70 %, 90 % and 100% for 5 mins each), embedded in Durcupan (44610-1EA, Sigma-Aldrich) and polymerised at 65 °C for 48 hrs. The glass coverslip was removed from the resin by submerging in liquid nitrogen, and the cells of interest relocated using the alphanumeric grid imprinted on the surface of the resin block. The block was trimmed to the cell of interest, serial sectioned at a thickness of 70 nm using a UC7 ultramicrotome (Leica Microsystems) and a diamond knife (Diatome) and collected onto 2 x 1 mm copper slot grids with a formvar support film (G089, TAAB Laboratories Equipment). Sections were post-stained using Reynold’s lead citrate and 1 % uranyl acetate. Serial images of the cell were then acquired using a Transmission Electron Microscope (TEM; Tecnai Spirit BioTwin; ThermoFischer Scientific) operated at 120 keV.

Serial TEM images were aligned using the TrakEM plugin in FIJI [65] (www.fiji.sc) and aligned to the confocal images using BigWarp (Bogovic et al., 2016) (www.fiji.sc).

### Correlative light and focused ion beam scanning electron microscopy

Cells were cultured, fixed and imaged using confocal microscopy as described above. For post-fixation and staining, a modified version of the NCMIR method was used (Deerinck, et al., 2010). Cells were post-fixed in 2.5 % glutaraldehyde/ 4 % formaldehyde in 0.1 M PB (pH 7.4) at RT for 30 mins, and stained in 1 % reduced osmium (1 % osmium/ 1.5 % potassium ferricyanide) at 4 °C for 1 hr. Cells were then treated with 1% thiocarbohydrazide (TCH) for 20 mins at RT, followed by 2 % osmium tetroxide for 30 mins at RT, and incubated overnight at 4°C in 1% uranyl acetate. The following day, cells were stained *en bloc* with lead aspartate (pH 5.5) at 60°C for 30 mins. The glass coverslip was then removed from the Mattek dish, and underwent a graded dehydration in ethanol (25 %, 50 %, 70 %, 90 % and 100 %, 5 mins per step) followed by embedding in Durcupan and polymerisation at 65 °C for 48 hours.

The coverslips were subsequently removed from the resin block using liquid nitrogen, and the cells relocated using grid coordinates. Areas of 3 x 3 grid squares in size with the cell of interest located at the center were then trimmed out using a razor blade, removed from the resin block (Russell, M.R.G et al, 2017), and mounted on a 12.7 mm SEM pin stub (10-002012-100, Labtech) using silver paint (AGG3691, Agar Scientific). After mounting, each sample was sputter coated with a 10 nm layer of platinum (Q150S, Quorum Technologies).

Focused ion beam scanning electron microscopy (FIB SEM) was carried out using a Crossbeam 540 FIB SEM with Atlas 5 for 3-dimensional tomography acquisition (Zeiss, Cambridge). The cells of interest were relocated by briefly imaging through the platinum coating at an accelerating voltage of 20 kV with 500 pA current, and correlating to the previously acquired confocal images. Once relocated, reoriented, and prepared for serial imaging, electron micrographs of the region of interest were acquired at 5 nm isotropic resolution, using a 11 µs dwell time. During acquisition, the SEM was operated at an accelerating voltage of 1.5 kV with 1 nA current. The EsB detector was used with a grid voltage of 1,200 V. Ion beam milling was performed at an accelerating voltage of 30 kV and current of 700 pA.

After cropping to the region of interest using Fiji [65], the datasets comprised 2,337 slices with an approximate volume of 1,695 µm^3^ (21.3 µ m x 6.8 µ m x 11.7 µ m) for sample 51, and 2,001 slices with an approximate volume of 1,428 µ m^3^ (17.2 µ m x 8.3 µm x 10 µm) for sample 52. Each stack was registered using the Atlas 5 autotune marks (template matching by normalized cross correlation; https://sites.google.com/site/qingzongtseng/template-matching-ij-plugin) and batch processed to suppress noise and enhance sharpness and contrast (i. Gaussian blur 0.8 pixel radius; ii. smart sharpening with highlights suppressed: radius 10 pixels, strength 60%, then radius 1.2 pixels, strength 150 %; iii. 8-bit greyscale conversion; Adobe Photoshop 2020). For reorientation in the image plane of the matching confocal datasets, each image stack was rotated and resliced perpendicular to the original coverslip plane, prior to alignment with the confocal dataset using BigWarp (Bogovic et al., 2016).

### ELISA assay

To determine the concentration of α-Syn monomer or oligomer, cell lysates were collected after treatment with either WT or A53T unlabeled monomers at various time points. The collected lysates were stored at −80°C until use. α-Syn oligomer was analyzed using Human Synuclein, alpha (non A4 component of amyloid precursor) oligomer (SNCA oligomer) ELISA kit (CSB-E18033h, Generon) and monomeric α-Syn was analyzed using LEGEND MAX™ Human α-Synuclein ELISA Kit (SIG-38974, BioLegend). Then it was normalized by total protein per well using Pierce BCA Protein Assay Kit (23225, Thermo Fisher Scientific).

### Single vesicle based membrane permeabilization assay

For membrane permeabilization assay, vesicles are prepared as previously described [35]. Using this assay, it has been previously shown that α-Syn oligomers disrupt and permeabilize membranes [66, 67]. Briefly, vesicles are synthesized using Phospholipids 16:0-18:1 PC and biotinylated lipids 18:1-12:0 Biotin PC using freeze thaw method with mean diameter of 200 nm. Each vesicle is filled with 100 μM Cal-520 dye and immobilized in PLL-g-PEG coated plasma cleaned glass coverslips using biotin-neutravidin linkage. The surroundings of the vesicles were filled with Ca^2+^ buffers. 50 µL of sample was incubated with the vesicles for 15 minutes and Ca^2+^ influx was quantitatively measured using a homebuilt Total Internal Reflection Fluorescence Microscope (TIRFM) based on an inverted Nikon Ti microscope. The 488 nm laser was a focused back-focal plane of the 60X, 1.49NA oil immersion objective lens used to excite the Cal-520 dye. The fluorescence signal was collected by the same objective and passed through an appropriate set of filters before imaged in EMCCD camera. To check if the aggregate present in SNCA media and lysates are composed of α-Syn, we used previously reported methods to determine the composition of the aggregates (De et al., 2019; Drews et al., 2017). Media was incubated with Anti-Alpha-Synuclein (phospho S129) antibody [EP1536Y] (Abcam ab51253) for 30 minutes and then added to the coverslips containing dye filled vesicles.

### Liposome preparation

Cardiolipin, 1,2-dimyristoyl-sn-glycero-3-phosphocholine (DMPC) (Avanti polar lipids) and biotinylated cardiolipin (Echelon Biosciences) were obtained as a lyophilized powder. About 1.5 mg of each lipid was transferred into glass test tubes and dissolved in 2 ml of chloroform: methanol (3:1) solvent. A thin layer of lipid was prepared by continuously rotating the test tube while treating the sample with a gentle stream of N_2_ gas until all the solvent evaporated. Test tubes were kept under vacuum overnight to remove any trace amount of solvent. The dry lipid layers were rehydrated with an appropriate amounts of 25 mM Tris buffer (pH 7.4) to make a final concentration of 2 mM and vortexed for 2 mins to dissolve the lipid layer into buffer. Biotinylated cardiolipin liposomes were prepared in a similar way by adding 2 % (M/M) biotin-conjugated cardiolipin before processing for liposomes preparation. Mixed vesicles followed an extra step of incubation at 40 °C with constant stirring to allow homogenous distribution of lipids in the liposomes. Small unilamellar vesicles were prepared by bath sonication (VWR ultrasonic) until a clear solution was obtained, indicating formation of unilamellar vesicles. Lipid vesicles were stored at 4 °C in glass vials until further use.

### Aggregation assay

ThT fluorescence time-course measurements were performed in 96-well microliter plates using FLUOstar Omega microplate reader (BMG Labtech) with excitation and emission wavelengths set to 450 nm and 485 nm. 50 µM protein samples were incubated in the presence and absence of lipid vesicles (400 µM) with 20 µM ThT in 25 mM Tris buffer (pH 7.4). Each sample-well contained 100 µL of the reaction mixture and spontaneous aggregation was induced by incubation at 37 °C without shaking. A minimum of 3 trails were performed to ensure its reproducibility.

### Circular dichroism spectroscopy (CD spectroscopy)

A Jasco J-810 spectrometer (Jasco) was used for circular dichroism (CD) measurements. Samples containing 10 µM WT α-Syn alone and in the presence of cardiolipin (80 and 160 µM) were prepared in 25 mM phosphate buffer (pH 7.4) and measured immediately after mixing. Experiments were performed at 25 °C using a quartz glass cell with 1 mm path length. The samples were recorded from 190 nm to 260 nm wavelength with a total of three scans for each sample.

### Transmission electron microscopy (TEM)

Selected fibril samples from 96 well plate was diluted to 3 µM in 25 mM phosphate buffer (pH 7.4). An aliquot (7 μL) of diluted sample was placed onto carbon support film 300 mesh, 3 mm copper grids (TAAB) and allowed for 3 min and blot dried. The TEM grids were subsequently stained using 7 μL of 1 % uranyl acetate for 3 min followed by blot drying. Samples were imaged on a Thermo Fisher Scientific Tecnai F20 electron microscope (200 kV, field emission gun) equipped with an 8k x 8k CMOS camera (TVIPS F816).

### Slide preparation for TIRF microscopy

Biotinylated liposomes were prepared as described previously [62]. Briefly, glass coverslips (22 × 22 mm) were cleaned via incubation in an argon plasma oven for 1h prior to use. Plasma cleaned coverslips were affixed to Frame-Seal slide chambers (9 × 9 mm^2^, Biorad, Hercules, CA) and the chamber was filled with about 50 µl poly-L-lysine solution (Sigma Aldrich) and incubated for 30 minutes at room temperature in a covered box. Glass coverslips were washed twice with filtered Tris buffer, after which sample containing α-Syn fibrils prepared in the presence of cardiolipin was added in the chamber, and the coverslip was placed on the microscope stage for imaging.

### TIRFM imaging

Imaging was performed in a home built total internal reflection microscope as described previously [68]. Amyloid fibrils prepared in the presence of biotin-conjugated cardiolipin were incubated with 1 nM Alexa Fluor 647 (AF647)-streptavidin (Thermo Fisher Scientific) and 5 mM ThT (Sigma Aldrich) for 10 minutes at room temperature. Images were recorded for 50 frames from the red channel (AF647 emission) with 641 nm illumination, followed by green channel (ThT emission) with 488 nm illumination. All the experiments were performed in two color coincidence detection (TCCD) mode.

### Immunohistochemistry

Cells were fixed in 4 % paraformaldehyde and permeabilized with 0.2 % Triton-X 100. 5 % BSA was used to block non-specific binding before cells were incubated with primary antibodies either for 2 hours at room temperature or overnight at 4 °C. The next day, cells were washed three times with PBS and incubated with secondary antibody for 1hr at room temperature. Cells were mounted with antifading medium after three times wash steps (DAPI was added in the second wash if required) and let dry overnight.

*Lists of primary antibodies used;* Anti-GFAP antibody (abcam, ab7260, 1:500), Anti-beta III Tubulin antibody (abcam, ab78078, 1:500), Anti-K67 antibody (abcam, ab254123, 1:500), Recombinant Anti-Alpha-synuclein aggregate antibody [MJFR-14-6-4-2] - Conformation-Specific (abcam, ab209538, 1:200). *Lists of secondary antibodies used;* Goat Anti-Chicken IgY H&L (Alexa Fluor® 488) (abcam, ab150169, 1:500), Goat Anti-Mouse IgG H&L (Alexa Fluor® 555) (abcam, ab150114, 1:500), Goat Anti-Rabbit IgG H&L (Alexa Fluor® 647) (abcam, ab150079, 1:500).

### Aptamer staining

For ATTO 425 labelled Aptamer staining, cells were permeabilized with 0.25 % Triton X-100 and blocked with 10 % normal goat serum (NGS) for 20 min followed by another 3 hours with 0.1 % Trion X-100 and 10 % NGS. Then cells were overnight incubated with 0.5uM Aptamer. After washing three times with PBS, cells were mounted with antifading medium [19].

### Statistical analysis

Origin 2020 (Microcal Software Inc., Northampton, MA) software was employed for the statistical analysis and exponential curve fitting. Unpaired two sample t-test or one-way ANOVA corrected with a Bonferroni and Tukey was used. Experimental data are shown as means ± standard error of the mean (SEM), and P value is set at 0.05. n = number of wells, if not stated otherwise. Sample sizes for experiments were selected to capture (1) technical variation including numbers of cell/field of view and coverslips and (2) biological variations including numbers of animal batch for primary co-culture and independent inductions and clones or patient line for hiPSC derived neurons. F-statistics was used to estimate variance within each group. All experiments were performed in a count balance manner and data were collected and analyzed without bias.

### Availability of data and resources

Imaging and tabular datasets, code, software, and all protocols used for this study have been deposited in relevant repositories, and described in Appendix 1.

## Supplementary figures

**Figure S1.**
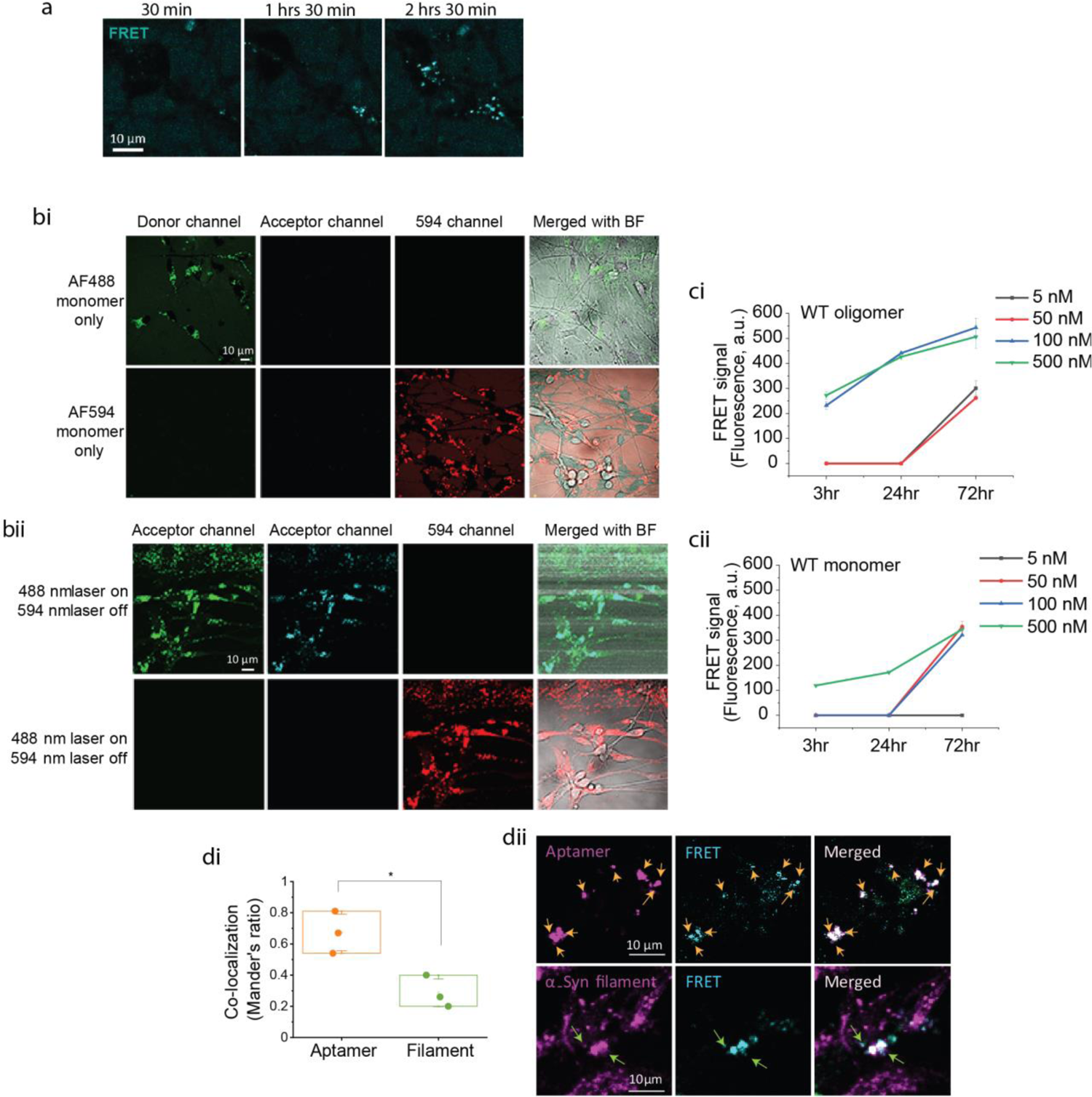
Related to main Figure 1. **(a)** Time-lapse images showing intracellular localization of α-Syn after incubation with α-Syn-AF488 and α-Syn-AF594 monomers. **(bi)** AF488-monomer or AF594-monomer alone does not induce FRET signal. **(bii)** FRET is not induced by 594nm excitation. **(ci & ii)** Intracellular FRET is concentration- and time-dependent. **(di & ii)** FRET fluorescence is co-localized with an amyloid oligomer-specific aptamer, and partially with an α-Syn filament antibody. The co-localization was quantified using Mander’s ratio.

**Figure S2.**
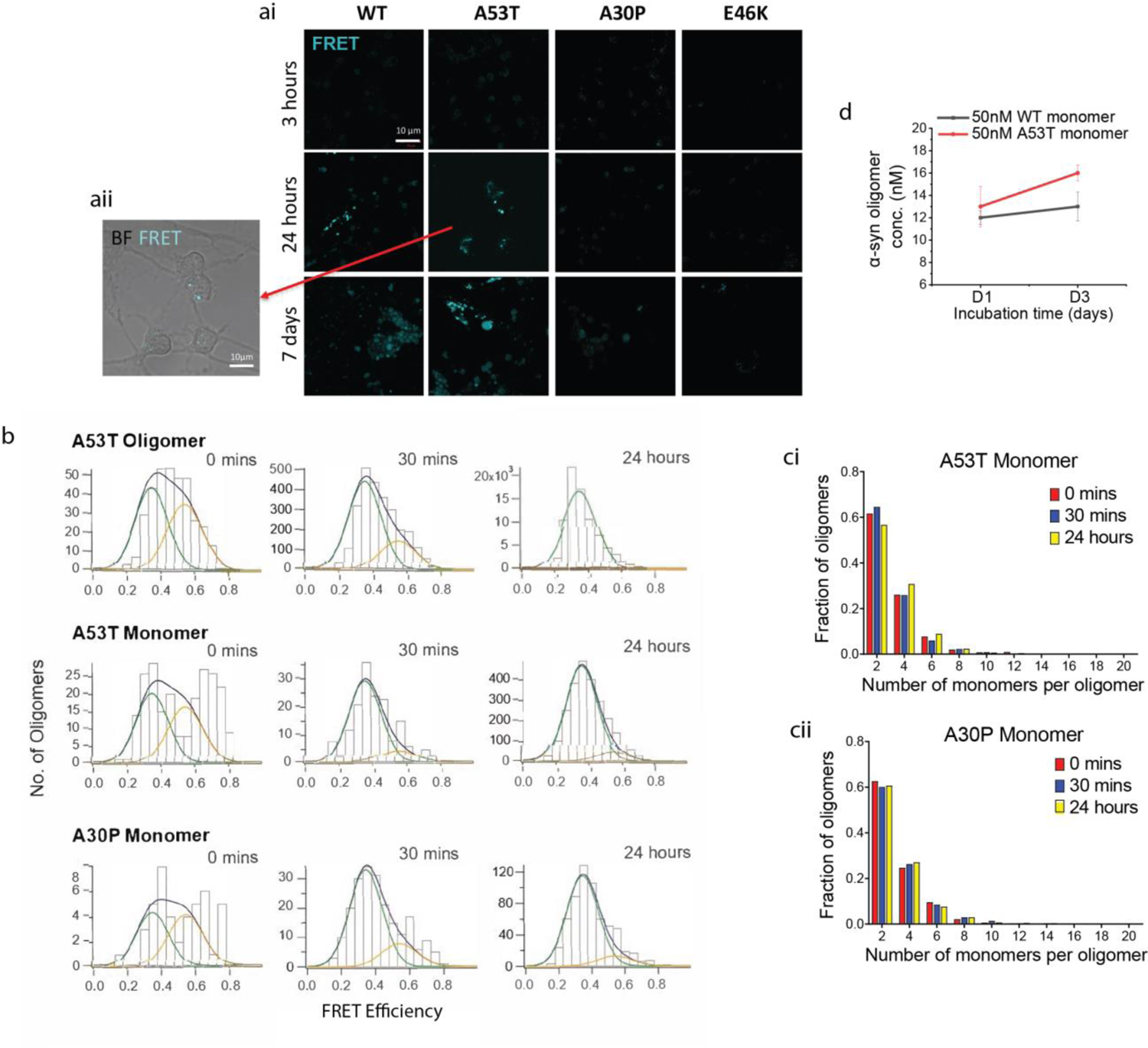
Related to main Figure 1. **(ai & ii)** Representative images of Figure 1D. **(b)** FRET efficiency histograms of lysates from Figure 1G. Oligomer populations were fitted (using a custom Igor script) for medium-sized oligomers (deemed to be between 3 and 20 subunits). Oligomers were determined to be either low FRET (green fit line) or high FRET (orange) based on their histogram population distributions at 24 hours, number of oligomers detected was highest for A53T oligomer, followed by A53T monomer, followed by A30P monomer. **(ci & ii)** Photobleaching data from A53T monomer and A30P monomer respectively, in parallel with data shown in Figure 1Hiii. **(d)** Higher oligomer concentration is detected in A53T monomer treated cells than WT after 3 days incubation using an α-Syn oligomer specific ELISA kit.

**Figure S3.**
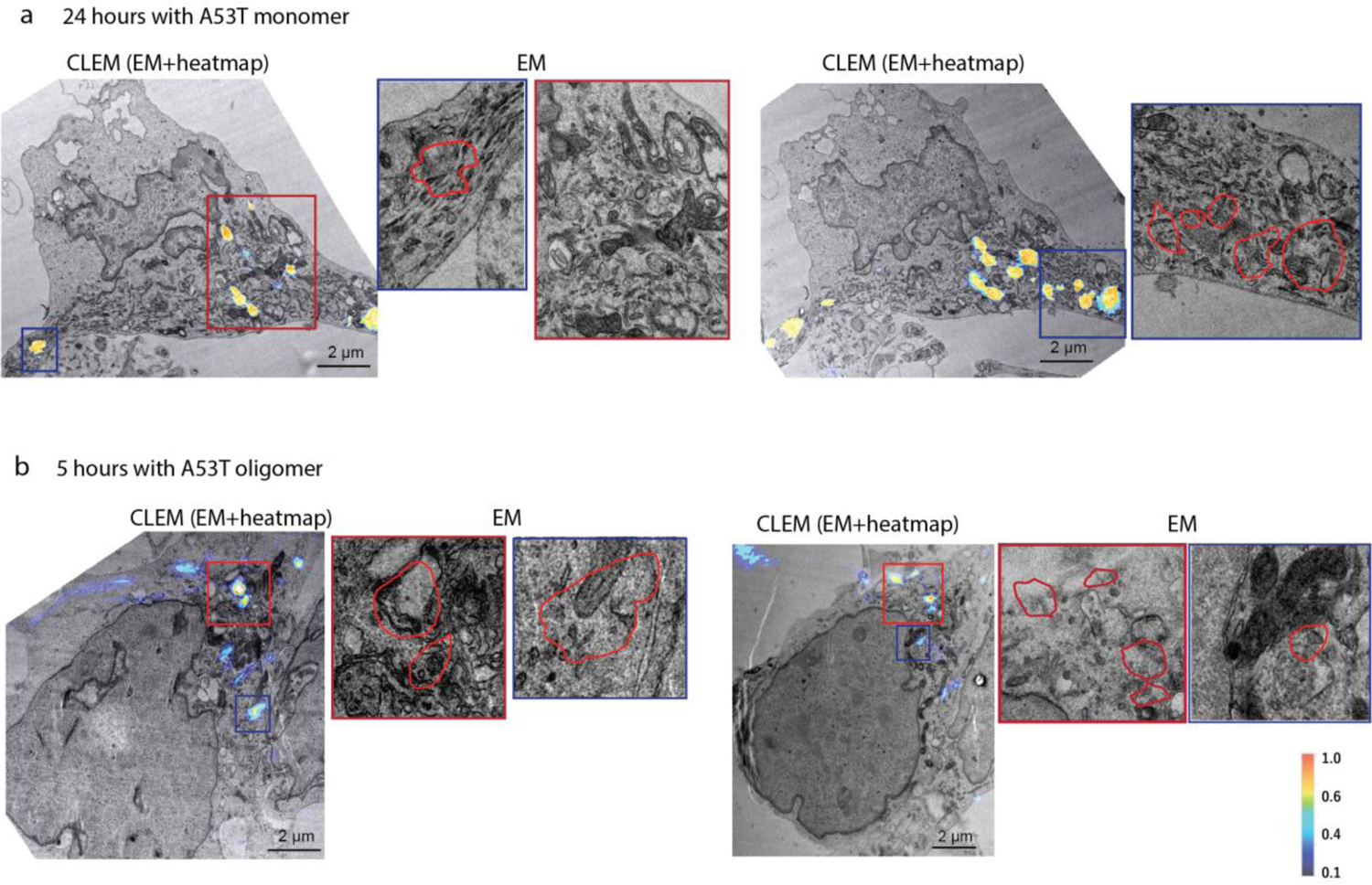
Related to main Figure 2. **(a & b)** FRET-CLEM images after addition of AF488 and AF594 labelled α-Syn at 24 hours (A53T monomer) and 5 hours (A53T oligomer) respectively. High (red) and low (blue) FRET areas are detected in human neurons.

**Figure S4.**
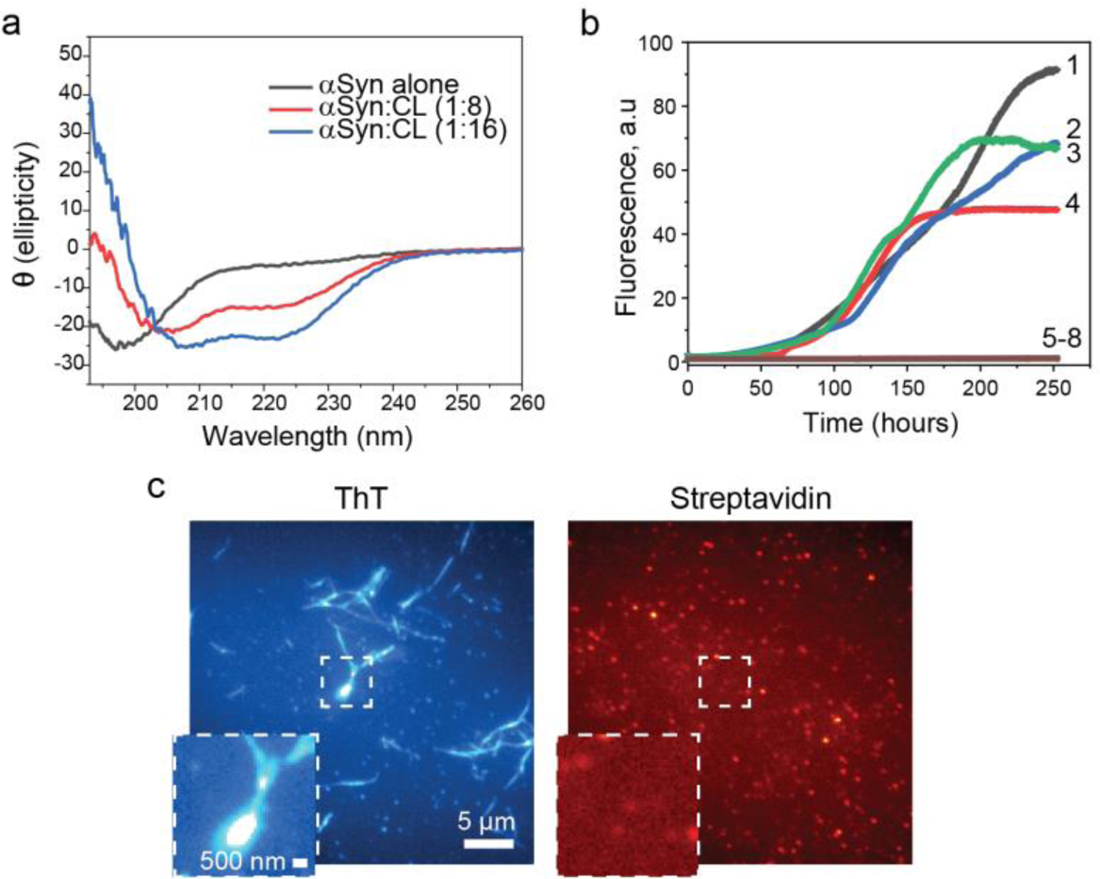
Related to main Figure 3. **(a)** Effect of cardiolipin on far-UV circular diagram (CD) spectra of α-Syn. Phosphate buffer, pH – 7.2, at 25 °C. In presence of cardiolipin, secondary structure of α-Syn changes from the random coil to alpha-helical form measured in pH 7.2 Phosphate buffer at 25 °C. In the absence of cardiolipin vesicles CD spectra was flat except minima at around 198 (black curve), which is characteristic spectra of a random coil. In the presence of CL vesicles at protein to lipid ratio 1:8 (red curve) and 1:16 (blue curve) a significant change occurred in the spectra, with minima at value at 222 nm and 208 nm, which is the characteristic spectra of the alpha-helical form of proteins. **(b)** Kinetics of amyloid formation by α-Syn (50 µM) in the presence of DMPS (curve 1 – 4) and in the presence of DMPC (curve 5 – 8). Measurements were performed at 30 °C and pH 7.4. Protein concentration was 50 µM and lipid vesicle concentration was 400 µM. ThT fluorescence was excited at 450 nm, and the emission wavelength was 482 nm. **(c)** Control images for Figure 3C.

**Figure S5.**
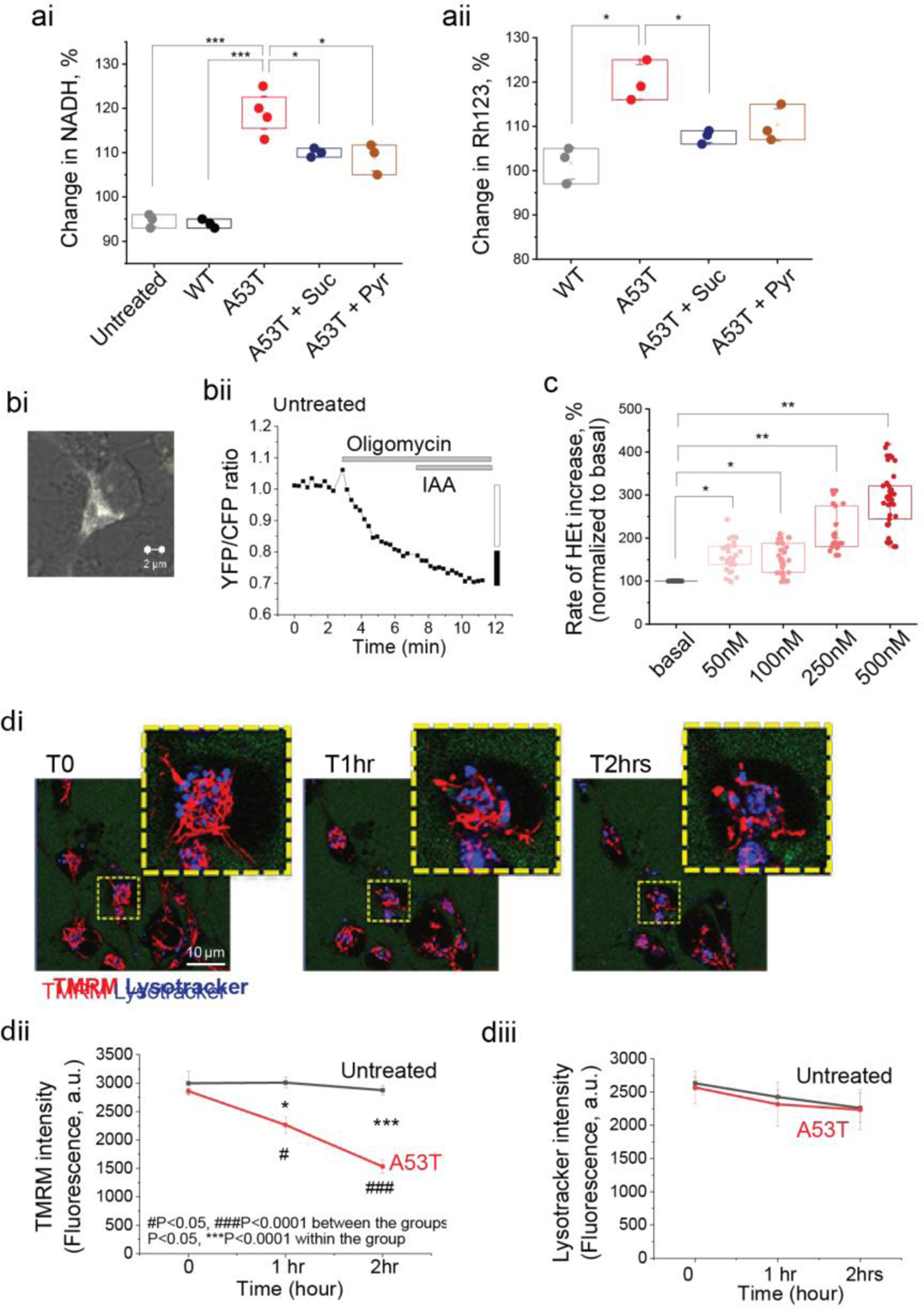
Related to main Figure 4. **(a)** Quantitative histogram referring to Figure 4A. **(bi)** Representative image of transfected cells with ATP probe. **(bii)** Representative trace of untreated cells in ATP measurement. **(c)** Superoxide is produced in a concentration dependent manner from α-Syn A53T monomer. **(di)** Time course images of mitochondria fragmentation after applying A53T α-Syn. **(dii & iii)** Δψ_m_ is significantly reduced after 2 hours whilst there is no change in lysosome after application of A53T α-Syn.

**Figure S6.**
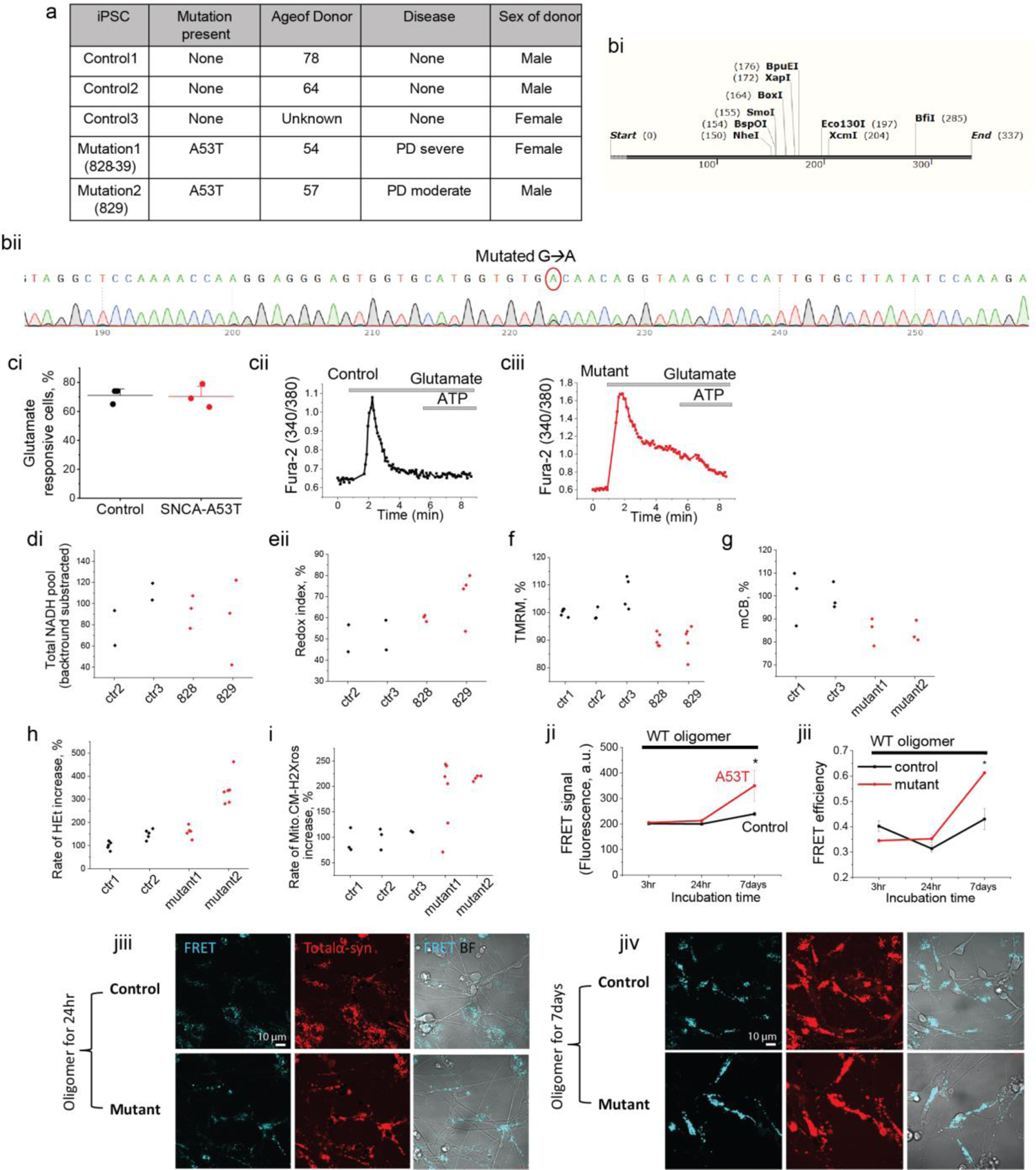
Related to main Figure 7. **(a)** Donor information of hiPSC used in this study. **(bi)** Design for Sanger sequence to confirm A53T point mutation on SNCA. **(bii)** The sequence confirms the presence of A53T mutation. **(ci – iii)** Assessment of neuronal function using [Ca^2+^] imaging. Both control and SNCA-A53T cells are responsive to 5 uM glutamate (control: 71.2 ± 3.1 %, A53T: 70.3 ± 4.6 %). **(d – i)** Separated data of each control (control 1, 2 and 3) and SNCA-A53T (828 and 829) lines referring to Figure 7. **(ji & ii)** Aftertreatment with α-Syn oligomer (1% of oligomer), neurons endogenously expressing SNCA-A53T mutation show higher FRET intensity and efficiency. **(jiii & iv)** Representative images showing increased FRET intensity after α-Syn oligomer treatment over time.

